# Integrative analysis from the epigenome through translation exposes patterns of dominant nuclear regulation during transient stress

**DOI:** 10.1101/479980

**Authors:** Travis A Lee, Julia Bailey-Serres

**Author notes:** Corresponding authors: Travis A Lee and Julia Bailey-Serres. T.L. is: Plant Biology Laboratory, The Salk Institute for Biological Studies, La Jolla, CA 92037, USA.

## Abstract

Gene regulation is modulated from chromatin to translation. To better understand the integration of nuclear and cytoplasmic gene regulatory dynamics, we performed a multi-omic survey of the epigenome through the translatome of the response of *Arabidopsis* seedlings to hypoxia and reoxygenation. This included eight assays of chromatin (histones, accessibility, RNAPII and transcription factor binding) and three assays of RNA (nuclear, polyadenylated, and ribosome-associated). Dynamic patterns of nuclear regulation distinguished stress-induced and growth-associated mRNAs. The rapid upregulation of hypoxia-responsive gene transcripts and their preferential translation was accompanied by increased chromatin accessibility, RNAPII engagement and reduced Histone 2A.Z association. The more progressive upregulation of heat stress gene transcripts was characterized by early engagement of RNAPII and elevation of nuclear over polyadenylated RNA. Promoters of the rapidly versus progressively upregulated gene cohorts were enriched for *cis*-elements of ethylene-responsive and heat shock factor transcription factor families, respectively. By contrast, genes associated with growth including ribosomal proteins underwent distinct histone modifications, yet retained RNAPII engagement and accumulated nuclear transcripts during the stress. Upon reaeration, many of the progressively upregulated and growth-associated gene transcripts were mobilized to ribosomes. Thus, multi-level nuclear regulation distinguishes transcript synthesis, accumulation and translation in response to a transient stress.

## Background

The regulated expression of protein coding genes in eukaryotes involves processes within the nucleus including remodeling of chromatin, recruitment of RNA polymerase II (RNAPII), co-transcriptional processing, and the export of mature mRNA to the cytoplasm, where it may be translated, sequestered and then degraded. These processes are modulated by conditions that necessitate alterations in metabolism and growth to maintain viability. In *Arabidopsis thaliana*, cellular hypoxia promotes activation of anaerobic metabolism to enable ATP production for basic cellular processes. This involves the rapid activation of transcription of 49 hypoxia responsive genes (*HRG*s) followed by the active translation of their mRNAs, while most other transcripts are sequestered from translation complexes until re-aeration (Branco-Price et al., 2005, 2008a; Juntawong et al., 2014; Sorenson and Bailey-Serres, 2014). The well translated *HRG* mRNAs lack notable sequences or features that can explain their effective translation (Branco-Price et al., 2005; Juntawong et al., 2014; Sorenson and Bailey-Serres, 2014), leading to the hypothesis that their preferential translational may reflect aspects of their nuclear regulation.

Transcriptional activation of many *HRG*s involves the evolutionarily conserved group VII ethylene response factor (ERFVII) transcription factors (TFs) that are required for survival of hypoxia (Xu et al., 2006; Hattori et al., 2009; Gibbs et al., 2011; Licausi et al., 2011; Yang et al., 2011; Abbas et al., 2015; Eysholdt-Derzsó and Sauter, 2017; Paul et al., 2016; Giuntoli and Perata, 2017). The five Arabidopsis ERFVIIs are stabilized under hypoxia due to attenuation of their oxygen-stimulated proteolysis via the Arginine branch of the N-degron pathway (Gibbs et al., 2011; Licausi et al., 2011; Paul et al., 2016; White et al., 2017; Millar et al., 2019). These include three constitutively synthesized ERFVIIs (RELATED TO APETALA 2.2, 2.3, and 2.12 [RAP2.2, RAP2.3, RAP2.12]) shown to transactivate *HRG* promoters in protoplasts through a hypoxia-responsive promoter *cis*-element (HRPE) identifiable in ∼50% of the *HRGs (Gasch et al., 2015; Giuntoli et al., 2014; Giuntoli and Perata, 2017)*. Two other ERFVIIs, HYPOXIA-RESPONSIVE ERF 1/2 (HRE1/2) are upregulated by hypoxia along with *ALCOHOL DEHYDROGENASE 1* (*ADH1*) and *PLANT CYSTEINE OXIDASE 1/2* (*PCO1/2*) that facilitate anaerobic metabolism and catalyze turnover of the ERFVIIs, respectively (Giuntoli et al., 2014, 2017; Weits et al., 2014; White et al., 2017).

Transcription requires binding of TFs to specific *cis*-elements typically located near the transcription start site (TSS) that facilitate assembly of an RNAPII initiation complex. The depletion of nucleosomes near a TSS is controlled by ATP-dependent chromatin remodelers and can be influenced by interactions with the transcriptional apparatus. TF binding and RNAPII activity both influence and are influenced by specific modifications and variants of histones(Talbert and Henikoff, 2017; Gates et al., 2017), which thereby provide a proxy of gene activity. For example, tri-methylated Histone 3-lysine 27 (H3K27me3) and H3K4me3 are prevalent in the gene body of lowly and actively transcribed genes, respectively (Asensi-Fabado et al., 2017; Gates et al., 2017). Acetylation of Histone H3 lysine residues (H3K9ac, H3K14ac) reduce interactions between DNA and histones and are associated with ongoing transcription(Gates et al., 2017), including genes actively transcribed during environmental stress in plants (Eberharter and Becker, 2002; Kim et al., 2008, 2012). A characteristic of heat-stress activated genes of Arabidopsis is high levels of the Histone 2A variant H2A.Z at the first nucleosome within the gene body, a feature proposed to reduce the energy required to commence transcriptional elongation at elevated temperatures (Kumar and Wigge, 2010; Sura et al., 2017; Cortijo et al., 2017; Dai et al., 2017; Torres and Deal, 2018).

As transcription commences, the phosphorylation of specific residues within the heptad repeats of the carboxyl terminal domain (CTD) of RNAPII orchestrate interactions with factors that facilitate transcription-coupled histone modifications as well as the co-transcriptional 5’ capping, splicing, and polyadenylation of the nascent transcript (Hajheidari et al., 2013; Milligan et al., 2016). CTD phosphorylation at Serine 2 (Ser2P) demarks active elongation (Phatnani and Greenleaf, 2006). In animals, pausing of RNAPII downstream of the TSS is common among genes activated by heat stress (Jonkers and Lis, 2015). The mapping of the 5’ end of nascent transcripts by Native Elongating Transcript sequencing indicated that elongation can be rate-limiting shortly after initiation in Arabidopsis seedlings (Zhu et al., 2018), although this was not discernable when nascent transcripts were surveyed by global nuclear run-on sequencing (GRO-seq) (Hetzel et al., 2016). Yet, both studies found transcription becomes rate-limited as RNAPII pauses just beyond the site of cleavage and polyadenylation. Following initiation, co-transcriptional intron splicing, polyadenylation and export are regulated by environmental cues including hypoxia (Branco-Price et al., 2008a; Mustroph et al., 2009; Juntawong et al., 2014; Van Veen et al., 2016; de Lorenzo et al., 2017; Niedojadło et al., 2016).

To better understand the integration of nuclear and cytoplasmic regulation of gene activity in response to hypoxia and reoxygenation we carried out a multi-omic study of dynamics in histones, open regions of chromatin, HRE2 binding and RNAPII-Ser2P distribution. Eight chromatin readouts were compared with modulation of nuclear RNA (nRNA), polyadenylated RNA (polyA RNA) and polyadenylated ribosome-associated RNA. The meta-analysis of these data exposed patterns of histone modifications, dynamic regulation of chromatin accessibility, *cis*-element enrichment, and temporal regulation of nRNA that accompanied dynamics in mRNA abundance and translation. This exposed underappreciated layers of gene regulation including rapid and complete upregulation of *HRGs*, progressive upregulation of heat stress-response genes, and incomplete downregulation of genes associated with growth, including ribosomal proteins. Both transcriptional elongation and nuclear export appear to be points of regulation in response to oxygen availability. The dataset provides a resource for evaluation of epigenetic state and RNAPII activity through translation in a model organism.

## Results

### A multiscale dataset for analysis of dynamics from chromatin to translation

Here, generated 11 distinct chromatin and RNA-based genome-scale datasets to evaluate gene regulatory dynamics in seedlings treated with normoxia (2h, 2NS; 9h, 9NS), non-lethal hypoxic stress (2HS, 9HS), and reaeration (2HS followed by 1-2 h NS, R). All growth and treatment conditions were consistent with prior analyses (Branco-Price et al., 2008a; Mustroph et al., 2009; Juntawong et al., 2014) (Figure 1a; Supplemental Figure 1a). Chromatin immunopurification and sequencing (ChIP-seq) was used to survey the position and abundance of H3K9ac, H3K14ac, H3K4me3, H3K27me3, and H2A.Z along genes. The capture of nuclei by Isolation of Nuclei Tagged in Specific Cell Types (INTACT) (Deal and Henikoff, 2010; Maher et al., 2018) was coupled with Assay for Transposase-Accessible Chromatin using sequencing (ATAC-seq) to map regions of chromatin depleted of nucleosomes (Buenrostro et al., 2013). We also used ChIP-seq to survey regions bound by RNAPII-Ser2P and the hypoxia-induced ERFVII HRE2. Finally, three transcript populations defined by methods of purification were assayed: nRNA (rRNA subtracted nuclear transcriptome obtained by INTACT (Reynoso et al., 2017)), polyA RNA selected by binding to oligo(dT) beads (transcriptome), and TRAP RNA obtained by Translating Ribosome Affinity Purification and oligo(dT) selection (translatome) (Juntawong et al., 2014; Zanetti et al., 2005). Each RNA population was reproducible and distinguishable when plotted by t-SNE (Figure 1b-c; Supplemental Figure 1).

**Figure 1.**
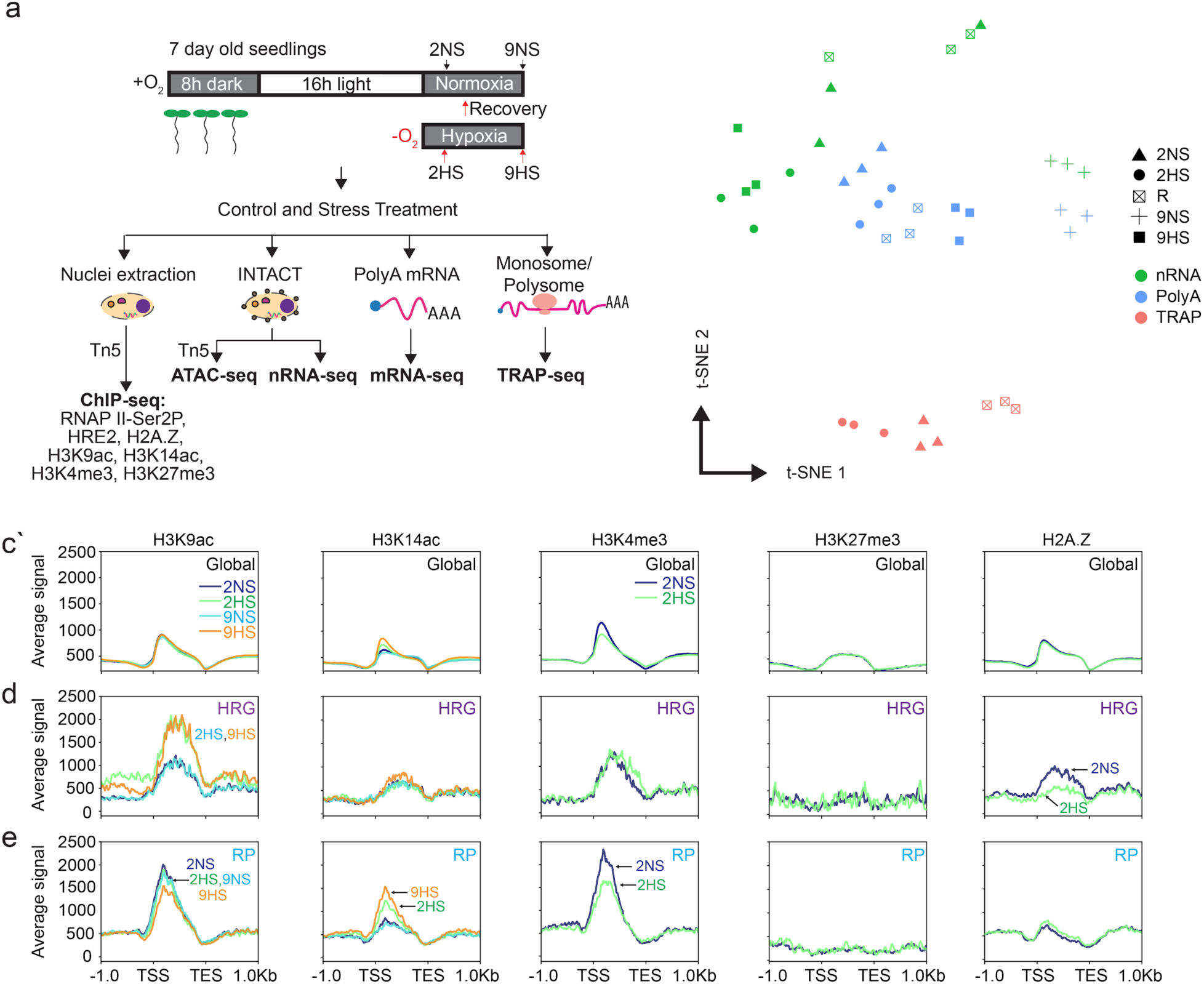
Multiscale chromatin and RNA gene regulatory analyses of control (normoxic) and hypoxic seedlings of *Arabidopsis*. **a** Schematic of experiments performed. Seven-day-old seedlings were subjected to control (normoxic) (2NS, 9NS), hypoxic stress (2HS, 9HS), or re-oxygenation following 2HS (R) conditions. Vertical arrows indicate time of harvest. Chromatin immunopurification (ChIP) was performed to evaluate genomic regions bound by the Ser2 phospho-isoform of RNA Polymerase II (Ser2P), the Histone 2 variant H2A.Z, modified Histone 3 (H3K4me3, H3K27me3, H3K9ac, H3K14ac), and the group VII Ethylene Responsive Factor (ERF) transcription factor HYPOXIA RESPONSIVE ERF 2 (HRE2). Isolation of Nuclei TAgged in specific Cell Types (INTACT) purified nuclei were used for Assay for Transposase Accessible Chromatin (ATAC) and purification of nuclear RNA. Ribosome-associated mRNA was obtained by Translating Ribosome Affinity Purification (TRAP). **b** Individual replicate samples of nuclear [nRNA], polyadenylated mRNA [polyA] and ribosome-associated polyA mRNA [TRAP]) were compared by t-Distributed stochastic neighbor embedding (t-SNE). **c-e** Distributions of histone modifications/variants across genic regions **c**, for the core hypoxia responsive genes (*HRG*s, *n*=49) **d,** and cytosolic ribosomal proteins (*RP*s, *n*=246) **e**. Read distributions are plotted from 1 kb upstream to 1 kb downstream of gene units defined by the transcription start site (TSS) and transcription end site (TES).

### Stress- and growth-associated genes contrast in chromatin accessibility, histone, modifications and RNA modulation under hypoxia

For the nucleosome level evaluation, the histone data were plotted along each annotated protein-coding gene and global averages were compared for each condition (Figure 1d; Supplemental Figure 2a). H3 modifications associated with transcription (H3K9ac, H3K14ac, H3K4me3) and stress-activated transcription (H2A.Z) were enriched just 3’ of the position of TSS and tapered off at the primary site of polyadenylation (TES), whereas H3K27me3 was distributed more evenly across gene bodies. Hypoxia had little effect on these modifications at the global level, with the exception of a slight elevation of H3K14ac and reduction of H3K4me3 near the TSS.

Dynamic alteration in histone modification, abundance, and distribution was observed for the induced and translated *HRG*s and stable but poorly translated cytosolic *RIBOSOMAL PROTEIN* (*RP*) genes. *HRGs* significantly increased in H3K9ac and decreased in H2A.Z association at 2HS, with limited changes in these marks on *RP*s (Figure 1e, f; Supplemental Figure 1b, S3a-c). H3K4me3 levels were notably reduced on the *RP*s but not *HRG*s in response to 2HS. When the stress was extended from 2 to 9 h, H3K9ac significantly increased on 863 and 3,646 genes and H3K14ac on 2 and 1,216 genes after 2HS and 9HS, respectively (Supplemental Supplemental Table 2). A progressive increase of H3K14ac but not H3K9ac on *RPs* was accompanied by little change in steady-state transcript abundance (Figure 1f; Supplemental Figure 1c; Supplemental Figure 3a,c). These data indicate that these two gene cohorts undergo distinct epigenetic regulation in response to hypoxia.

Next, we evaluated dynamics in chromatin accessibility in genic regions by ATAC-seq (Figure 2a-b). Accessibility in 5’ promoter regions is associated with depletion of nucleosomes and binding of TFs and other transcriptional machinery (Lu et al., 2016; Sijacic et al., 2017; Maher et al., 2018), whereas accessibility in 3’ flanking regions is associated with formation of chromatin loops and transcriptional termination (Ansari and Hampsey, 2005; Ozsolak et al., 2008). The ATAC-seq reads mapped almost exclusively within 1000 bp 5’ of the TSS and ∼300 bp 3’ of the major TES of genes in all four conditions (Figure 2c). The average ATAC signal increased within the promoters of both *HRGs* and *RP*s (1.8- and 1.4-fold, respectively) after 2HS, yet ATAC reads showed expansion into more 5’ and 3’ flanking regions on the *HRG*s. Peak-calling of Tn5 hypersensitive insertion sites (THSs) across the genome identified constitutively present (8,072) and and an even larger group of stress-specific THSs (25,795) (Figure 2d-f; Supplemental Table 1a).

**Figure 2.**
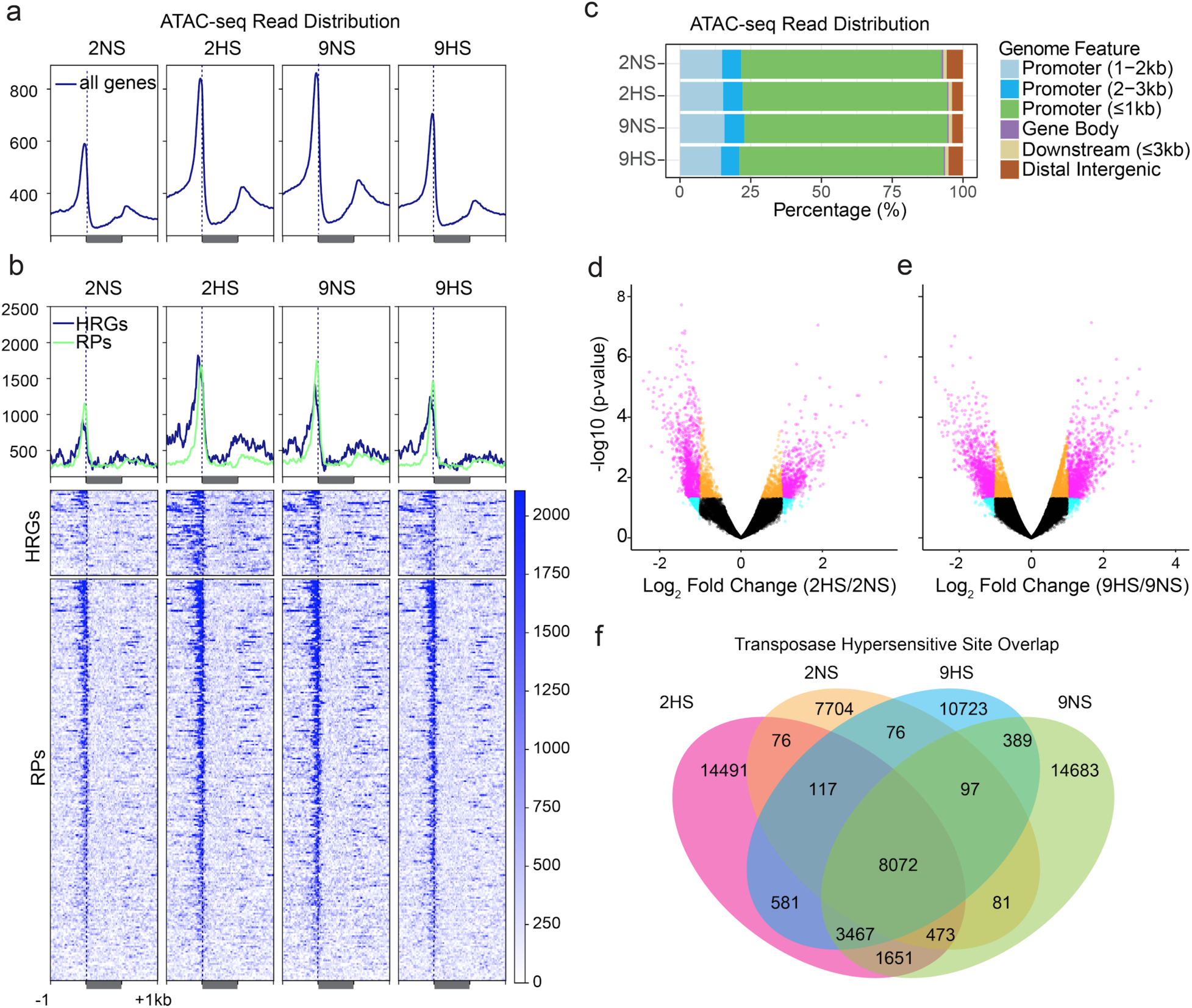
Regions of chromatin accessibility determined by Assay of Transposase-Accessible Chromatin using sequencing (ATAC-seq) are dynamically regulated by hypoxic stress. **a-b** Average ATAC-seq read distribution of chromatin accessibility for protein-coding genes (**a**), HRGs and RPs (**b**). **c** Distribution of transposase hypersensitive sites (THSs) on genomic features for each condition. **d-e** Volcano plot of the log_2_ fold change in THSs identifiable under two conditions and the significance value of their difference. Genes indicated in pink meet the criteria of log_2_ fold change value > |1|, 0.05 < p. value. **f** Overlap in THSs identified for each of four conditions.

To complete our survey of gene activity, we compared the distribution of the log_2_ fold change (FC) (2HS/2NS) values of the chromatin, RNAPII-Ser2P, and the four RNA populations at the genome-scale as well as for the *HRG*s and *RP*s (Figure 3a; Supplemental Table 2a). We found that stress-induced RNAPII-Ser2P engagement was generally correlated with increased H3K9ac (global R = 0.43; *HRGs* R = 0.17) and H2A.Z eviction (global R = −0.27), particularly for the *HRGs* (R = −0.79) (Figure 3 b,c; Supplemental Figures 3b, 4). *RPs* contrasted to the *HRGs*, displaying only minor changes in RNAPII-Ser2P association at 2HS but increased H3K14ac and nRNA elevation at 9HS (Figure 3a; Supplemental Figure 3b, c). These results further demonstrate that the epigenetic regulation of stress-induced and growth-associated genes is distinct under hypoxia.

**Figure 3.**
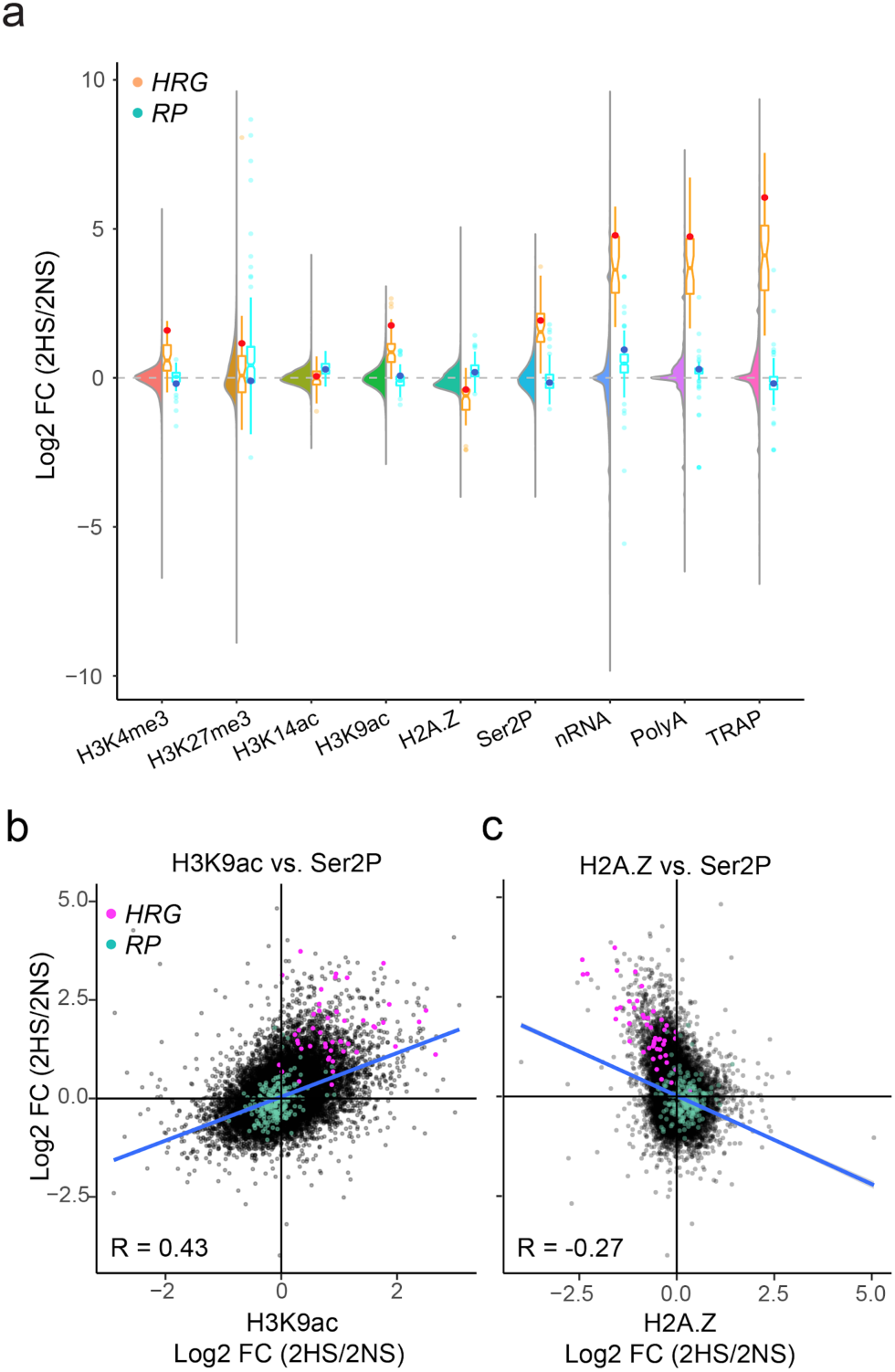
Short term hypoxic stress promotes pronounced changes in chromatin, transcription and nascent transcripts that are not uniformly reflected in transcript steady-state abundance or translation. Comparison of genome-wide data for chromatin and RNA for 2 h hypoxic stress (2HS) relative to the normoxic control (2NS). **a** Combined violin and boxplot of log_2_ fold change (2HS/2NS) of histone, RNAPII-Ser2P and RNA outputs. Violin, all genes; boxplot for the *HRG*s plotted in orange with *ALCOHOL DEHYDROGENASE 1* (*ADH1*) depicted as a red dot and the cytosolic *RP*s plotted in blue, with *RIBOSOMAL PROTEIN 37B* depicted as a dark blue dot. **b** Comparison of log_2_ fold change (2HS/2NS) between H2A.Z on the gene body and Ser2P on the gene body. **c** Comparison of log_2_ fold change (FC) (2HS/2NS) between H3K9ac and Ser2P on the gene body. For b and c, *HRG*s are plotted in pink and *RP*s in blue; Pearson’s coefficient of determination is shown for all genes.

### Regulation at nuclear and cytoplasmic scales can be concordant or discordant

Next we evaluated the generally presumed but untested hypothesis that *HRGs* are coordinately transcribed and translated under hypoxia. To do so, we performed a global-scale systematic analysis of the RNAPII-Ser2P, nRNA, polyA, and TRAP RNA datasets using co-regulation and cluster analyses. When we examined whether significantly up- and downregulated genes (DRGs; |log2 fold change| > 1; FDR < 0.05) were similarly regulated in all readouts of gene activity, we found that the previously non-surveyed nRNA had the greatest (1,722) and polyA RNA the fewest (602) up-DRGs (Figure 4a). The vast majority (92%) of the polyA up-DRGs were also upregulated in at least one other readout, but only 213 genes were significantly upregulated in all four readouts. Notably, these coordinately up-DRGs included 38 of the 49 *HRG*s. A parallel analysis found 72% of the polyA down-DRGs were significantly reduced in at least one other dataset with only 10 coordinately down-DRGs (Figure 4b). nRNA displayed the greatest number of down-DRGs (Ser2P [206], nRNA [2,608], polyA [291], TRAP [578]).

**Figure 4.**
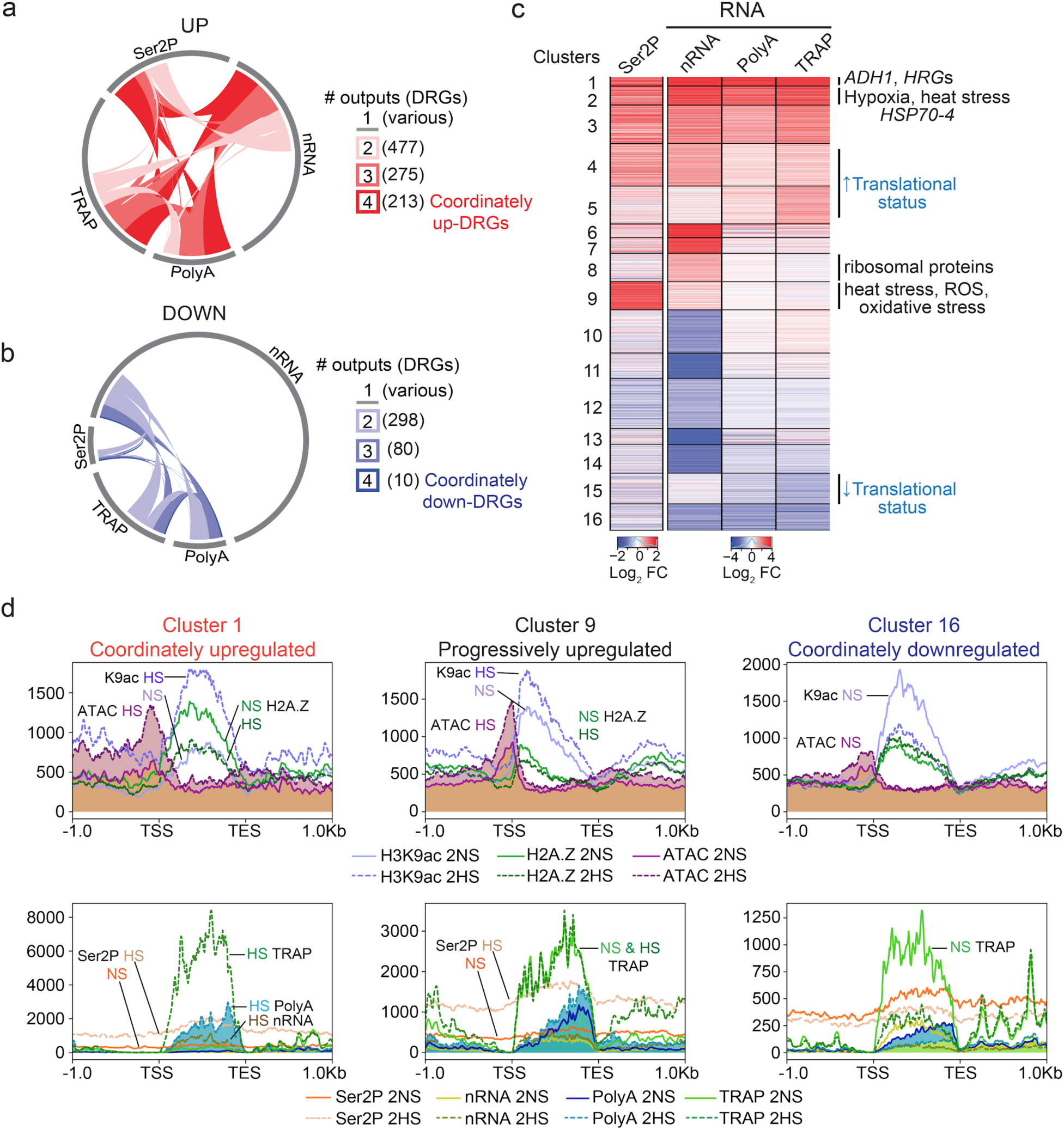
Multi-scale analysis of hypoxia-regulated genes reveals concordant and discordant patterns of gene regulation. Number of **a** up- and **b** down-differentially regulated genes (DRGs) in response to 2 h of hypoxic stress (|log2 fold change (FC)| > 1; False Discovery Rate (FDR) < 0.05) identified in each of the four readouts related to RNA: Ser2P ChIP, nRNA, polyA RNA, and TRAP RNA. Arc of circle circumference indicated by the grey line represents the DRGs in each of the four readouts. Genes differentially regulated in 2 to 4 readouts were tabulated (number in parentheses) and depicted by links within the circle. Genes differentially regulated in only one readout are represented by unlinked grey lines. **c** Heatmap showing similarly regulated genes based on four assays of gene activity. Partitioning around medoids clustering of 3,042 DRGs; 16 clusters. Selected enriched Gene Ontology terms are shown at right (p adj. < 1.47e-09), |FC| > 1, FDR < 0.05. Clusters displaying increased or decreased translational status (ribosome-associated [TRAP] relative to total abundance [polyA]) are shown. **d** Average signal of various chromatin and RNA outputs for genes of three clusters plotted from 1 kb upstream of the transcription start site (TSS) to 1 kb downstream of the transcript end site (TES) for the 2 h (2NS, 2HS) timepoints. Signal scale of graphs differ. Dashed lines used to plot hypoxic stress data. Upper panel, ATAC-seq data shaded. Lower panel 2HS polyA data shaded.

A lack of consensus in nRNA and polyA dynamics in response to the stress was not unexpected, as the nuclear RNA purification method did not require the presence of a polyA tail and therefore includes nascent transcripts. Indeed, the pairwise comparison of nRNA and polyA RNA abundance in the 2NS and 2HS samples identified transcripts that were enriched in nRNA or polyA RNA (Supplemental Figure 5; Supplemental Table 3). In both conditions, transcripts that were overrepresented in the nucleus included many associated with the cell cycle and gene silencing, whereas transcripts overrepresented in the polyA pool were associated with plastid and ribosome biogenesis and function. Our data reveal a complex nuclear and cytoplasmic regulatory landscape that was further explored by more integrative analyses of the chromatin and RNA datasets.

### Coordinate and progressive upregulation are characterized by distinct patterns of histone alterations

To better resolve genes with similar patterns of regulation, DRGs identified in at least one of the four datasets (n=3,042) were clustered into 16 groups and evaluated by Gene Ontology (GO) term enrichment (Figure 4c; Supplemental Table 4a). This confirmed coordinately up-DRGs (clusters 1-3) and down-DRGs (cluster 16). Genes in the remaining clusters that showed discordant regulation in one or more readout.

To better understand the regulatory variation at the chromatin, RNAPII, and RNA scales, we plotted the average signal value for each data type for each cluster along genic regions (Figure 4d; Supplemental Figure 6). This revealed that the rapid and complete upregulation of cluster 1-3 genes at 2HS was accompanied by enhanced chromatin accessibility, eviction of H2A.Z, as well as elevation of H3K9ac and RNAPII-Ser2P across the gene body. The concomitant rise in TRAP RNA indicated these transcripts were translated in proportion to their increased abundance, as shown for the *HRG*s by high-resolution ribosome footprint analysis (Juntawong et al., 2014). Not unexpectedly, the first two clusters were enriched for genes associated with decreased oxygen (cluster 1, p-adj. <6.73e^-22^) and general stress (cluster 2, <1.06e^-21^) (Supplemental Table 4a). Genome browser views of *ADH1* and *PCO2* illustrate the coordinate upregulation of *HRGs* in all four of the assays of gene activity (Figure 5a,b).

**Figure 5.**
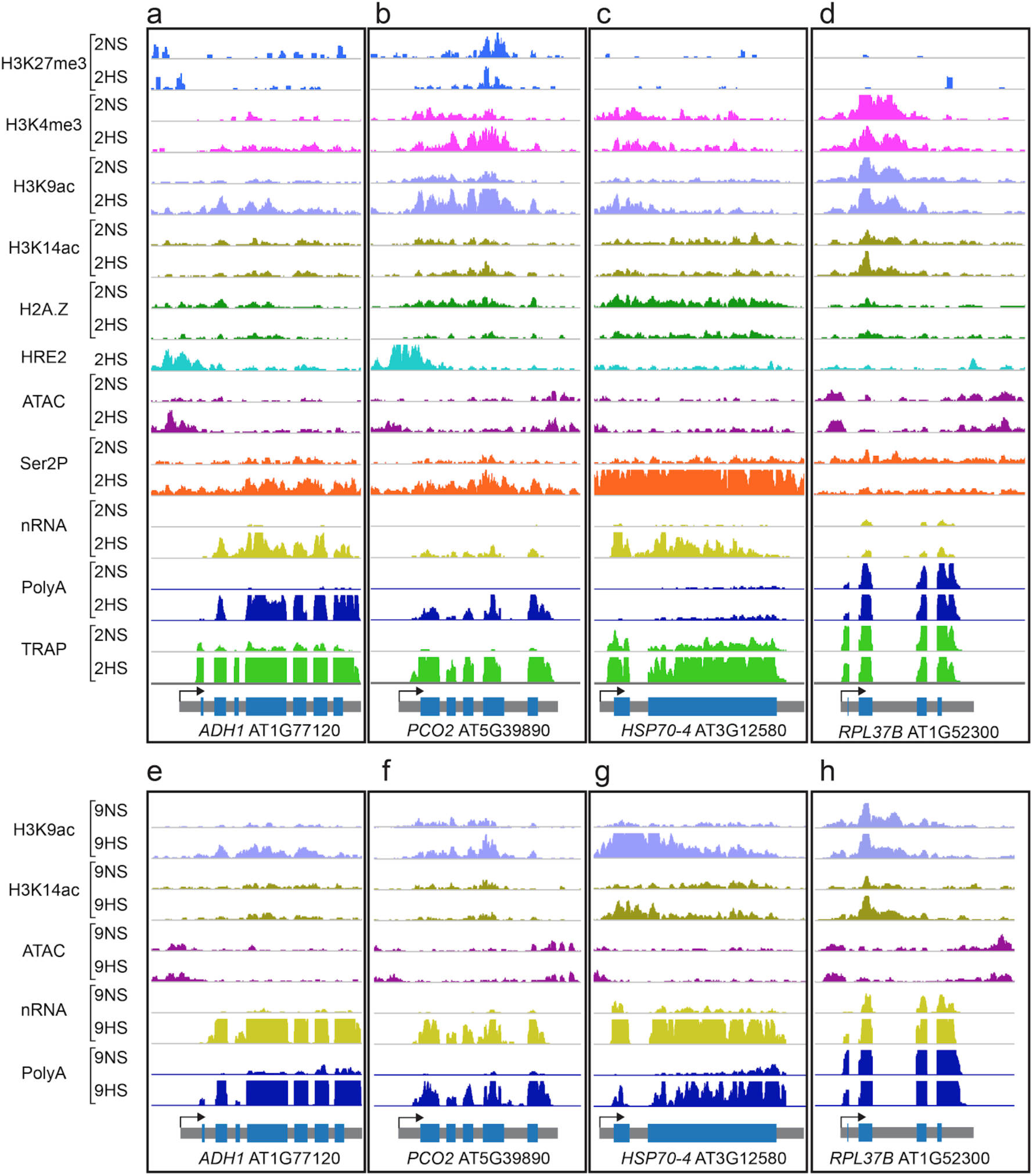
Genome browser view of normalized read coverage of histone, ATAC-seq, RNAPII-Ser2P, HRE2-chromatin immunopurification and RNA outputs for representative genes. **a-d** Normoxia (2NS) and 2 h hypoxic stress (2HS). **e-h** Normoxia (9NS) and 9 h hypoxia stress (9HS) samples. **a,e** *ALCOHOL DEHYDROGENASE 1* (*ADH1*), **b,f** *PLANT CYSTEINE OXIDASE 2* (*PCO2*), **c,g** *HEAT SHOCK PROTEIN 70-4* (*HSP70-4*) and **d,h** *RIBOSOMAL PROTEIN L37B* (*RPL37B*). The maximum read scale value used for chromatin based and RNA based outputs is equivalent for all genes depicted. At bottom, the transcription unit is shown in grey with the transcription start site marked with an arrow. *PCO2* shows a stronger decline in H2A.Z than *ADH1* in response to hypoxia at 2HS (*PCO2*, −1.15 log_2_ FC; *ADH1*, −0.33 log_2_ FC) and provides an example of elevated H3K4me3 under hypoxic stress.

Genes of clusters 4-8 were not upregulated to the same extent from RNAPII engagement through translation, indicating they are regulated by multiple mechanisms that are influenced by hypoxia. For cluster 4, the rise in nRNA coincided with elevation of H3K9ac and RNAPII-Ser2P, indicative of increased transcription, yet this was not accompanied by a concomitant rise in polyA or TRAP RNA (Figure 4c; Supplemental Figure 6). This incomplete upregulation could reflect nuclear retention or cytoplasmic destabilization of these transcripts. Clusters 6-8 genes showed elevated nRNA at 2HS, suggesting these transcripts are nascent or remain in the nucleus. Of these, cluster 8 displayed little change in H3K9ac but a rise in H3K14ac as well as nRNA at 2HS and 9HS (Supplemental Figures 6 and 7), consistent with the regulation of the *RP*s that are enriched in this cluster (Figure 3; Supplemental Figure 3b,c). This pattern of regulation is illustrated by *RPL37B* (Figure 5d).

One of the most distinct gene regulatory patterns was observed for cluster 9, which showed a dramatic increase in RNAPII-Ser2P engagement that was not accompanied by a rise in nRNA or polyA RNA. This could reflect an increase in RNAPII engagement (initiation) without completion of transcript synthesis or elevated rates of transcription accompanied by high cytoplasmic turnover. By comparison to cluster 1 genes, those of cluster 9 had lower and more 5’ skewed H2A.Z at 2NS that was only slightly reduced at 2HS (Figure 4d; Supplemental Figure 9a). The association of H3K4me3, a mark typical of actively transcribed genes, was significantly lower than genes of cluster 1 at 2HS (Supplemental Figure 9c,d). By 9HS the promoter accessibility, H3K9ac, H3K14ac, nRNA and polyA RNA abundance of cluster 9 genes increased (Supplemental Figure 7), indicating that they are progressively activated by hypoxia. Cluster 9 genes were enriched for heat (<6.20e^-13^) and oxidative stress (<8.23e^-11^), which were also prevalent in cluster 2 (heat [7.74e^-14^]; oxidative stress [<1.13e^-12^]). Cluster 2 was distinguishable from cluster 9 by induced Ser2P engagement that produced nRNA and translated RNA. *HEAT SHOCK PROTEIN 70-4* (*HSP70-4*), as an example, displayed a 3.4-fold increase in RNAPII-Ser2P across the gene body region accompanied with increased nRNA and TRAP RNA, but a delayed rise in polyA RNA (Figure 5c). The pronounced early engagement of RNAPII-Ser2P followed by delayed elevation of polyA RNA indicates genes of cluster 9 are progressively upregulated by hypoxia.

### Highly downregulated genes undergo distinct stimulus-regulated histone modifications

A survey of the down-DRG genes (clusters 10-16) demonstrated stimulus-driven dynamics of chromatin, RNAPII, and RNAs with two dominant patterns: coordinate downregulation (cluster 16) and incomplete downregulation (clusters 10-15) (Figure 4a). Cluster 10-14 were primarily distinguished by a significant reduction in nRNA at 2HS that was generally accompanied by increased H2A.Z and H3K14ac, and decreased H3K9ac downstream of the TSS (Supplemental Figures 6 and 7). Cluster 15 and 16 genes stood out as strongly and coordinately reduced in RNAPII-Ser2P engagement and at least two of the three RNA readouts at 2HS (Figure 4a; Supplemental Figure 6). These clusters displayed reduced chromatin accessibility, elevation of H2A.Z, and a sharp decline in H3K9ac,all of which were antithetical to the changes observed for the continuously up-DRGs (Supplemental Figure 9a,c). Clusters 15 and 16 were enriched for GO categories including: root development (cluster 15, <3.7e^-8^) and regulation of transcription and RNA biosynthesis (cluster 16, <7.78e^-7^). Despite their opposing regulation, the most highly up-(cluster 1-2) and down-DRGs (cluster 15-16) were characterized by H3 modifications that were more broadly distributed along the gene body than in other clusters. Thus, H2A.Z presence across a gene body, limited chromatin accessibility near the TSS, and reduced RNAPII engagement are signatures of restricted transcription under hypoxia.

### Prolonged hypoxia promotes variations of nuclear regulation of growth-associated genes

As seedling response to sub-lethal hypoxic stress changes over time (Branco-Price et al., 2008b), we compared the change in H3K9ac, H3K14ac, nRNA, and polyA RNA at 2HS and 9HS relative to untreated seedlings (Figure 6a,b; Supplemental Figure 8; Supplemental Table 2a). H3K9ac and H3K14ac levels were progressively yet distinctly altered for the *HRG* and *RP* cohorts (Figure 6a). Over 350 genes were progressively upregulated, displaying a rise in H3K9ac, nRNA, and polyA RNA at 2HS and 9HS (clusters 1b-2b; 38 *HRG*s). The remainder were incompletely up- or downregulated (clusters 4b-11b), indicating the regulatory patterns of genes within these clusters changed as hypoxia was prolonged. Many genes with elevated nRNA at 9HS were enriched for H3K14ac near their TSS (clusters 2b-8b) (Supplemental Figure 8), as exemplified by the *RPs* (enriched in clusters 5b and 8b). The increase in nRNA and H3K14ac without an accompanying rise in polyA RNA suggests that these genes are transcriptionally active but the nascent transcript remains in the nucleus or the mature (polyA) transcript is rapidly degraded. Other genes demonstrated dynamic up-(cluster 6b) or down-regulation between 2HS and 9HS (clusters 8b-11b). The downregulation at 9HS was pronounced for photosynthesis (cluster 8b, <5.8e^-7^ and <1.5e^-6^), root development (cluster 10b, <1.7e^-6^), and transcription (cluster 11b, <7.6e^-6^) (Supplemental Table 4b). A key finding was that temporal regulation of nuclear and polyA transcript abundance during hypoxia can be distinct.

**Figure 6.**
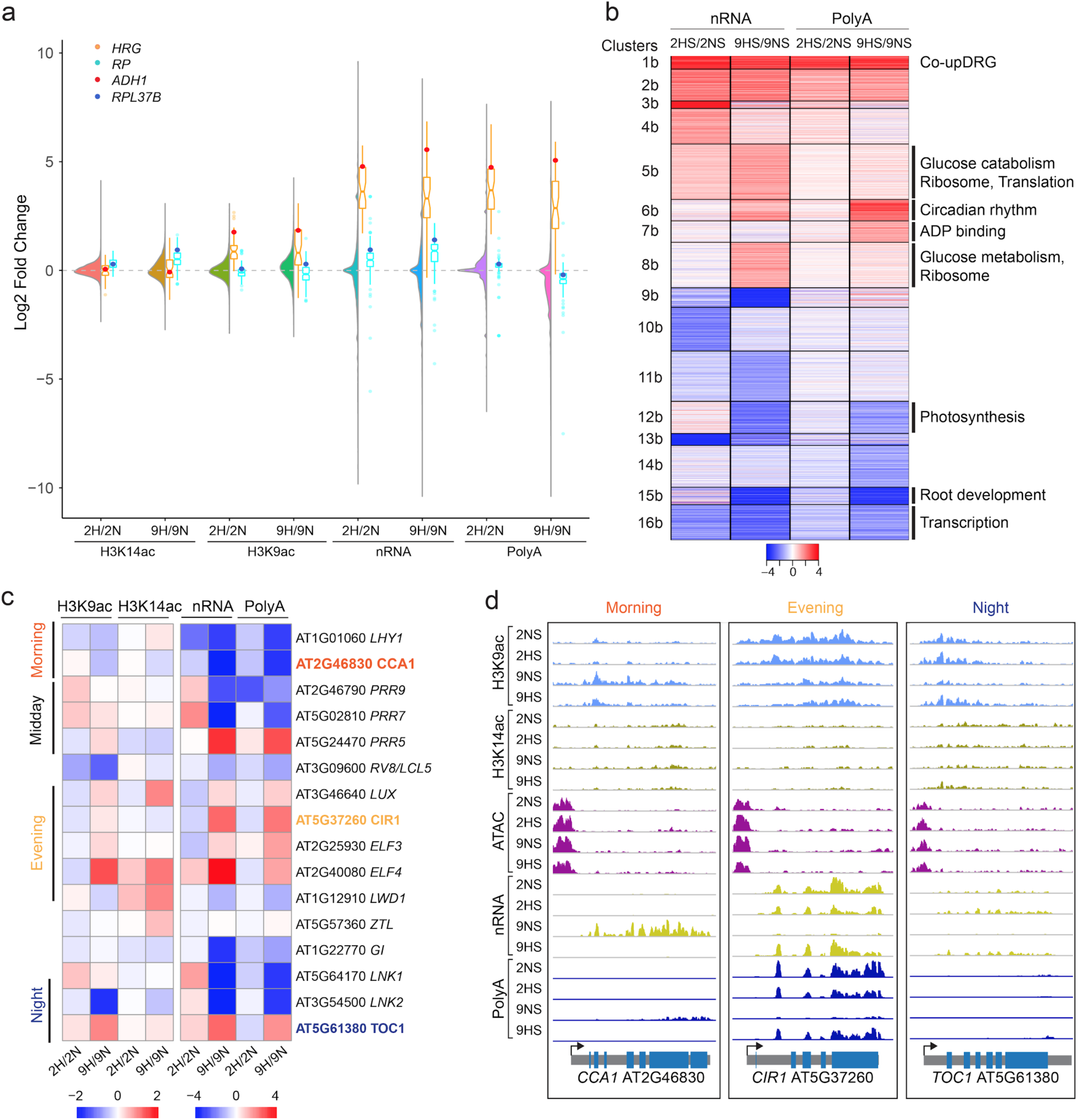
Prolonged hypoxia leads to further variation in epigenetic, transcriptional and posttranscriptional regulation. **a** Combined violin and boxplot of H3K9ac and H3K14ac, nRNA, and polyA RNA data. Violin plots of log_2_ fold change (9HS/9NS) data for all genes. Boxplots are data for the 49 *HRGs* plotted in orange with *ALCOHOL DEHYDROGENASE 1* (*ADH1*) depicted as a red dot and data for the cytosolic ribosomal proteins (RPs) plotted in blue with *RIBOSOMAL PROTEIN 37B* depicted as a dark blue dot. **b** Heatmap showing similarly regulated genes based on two dynamics in nRNA and polyA RNA. Partitioning around medoids clustering of 3,897 differentially regulated genes; 16 clusters. Selected enriched Gene Ontology terms are shown at right (p adj. < 1.37E-06), |FC| > 1, FDR < 0.05. **c** Heatmap of chromatin and RNA readouts of select circadian regulated genes under short and prolonged hypoxic stress. Heatmap scales between chromatin and RNA based readouts differ. **d** Brower views of a morning (*CIRCADIAN CLOCK ASSOCIATED 1*, CCA1), evening (*CIRCADIAN 1*, CIR1), and night (*TIMING OF CAB EXPRESSION 1*, TOC1) expressed genes. RNA scale of *TOC1* is three-fold lower than the scale used for *CCA1* and *CIR1*.

A number of genes associated with phasing of the circadian clock were perturbed when hypoxia was prolonged (cluster 6b; circadian rhythm <7.9e^-10^). Upon closer examination we found this included dampened upregulation of several morning gene and prolonged elevation of evening gene mRNAs, which normally peak towards the end of the light cycle when the stress was initiated (Figure 6c). Examples of this perturbation include the morning-expressed clock regulators *CIRCADIAN CLOCK ASSOCIATED 1* (cluster 16b) and *LATE ELONGATED HYPOCOTYL* (cluster 15b) which had dampened mRNA accumulation, and *TIMING OF CAB EXPRESSION* that had elevated polyA RNA after 9HS as compared to 9NS (Figure 6c,d). Thus, hypoxia imposed at the end of the light cycle delays or arrests circadian phasing.

### Many coordinately hypoxia-upregulated genes are targets of ERFVII regulation

We hypothesized that the coordinately up-DRGs might be transcriptionally activated by the low-oxygen stabilized ERFVIIs. Motif scans for the HRPE within promoters (500 bp 5’ of the TSS) across the genome confirmed significant enrichment of this *cis*-motif in the cluster 1 genes resolved in the 2HS analysis (Figure 7a; Supplemental Table 5), including the genes with strong coordinate upregulation.

**Figure 7.**
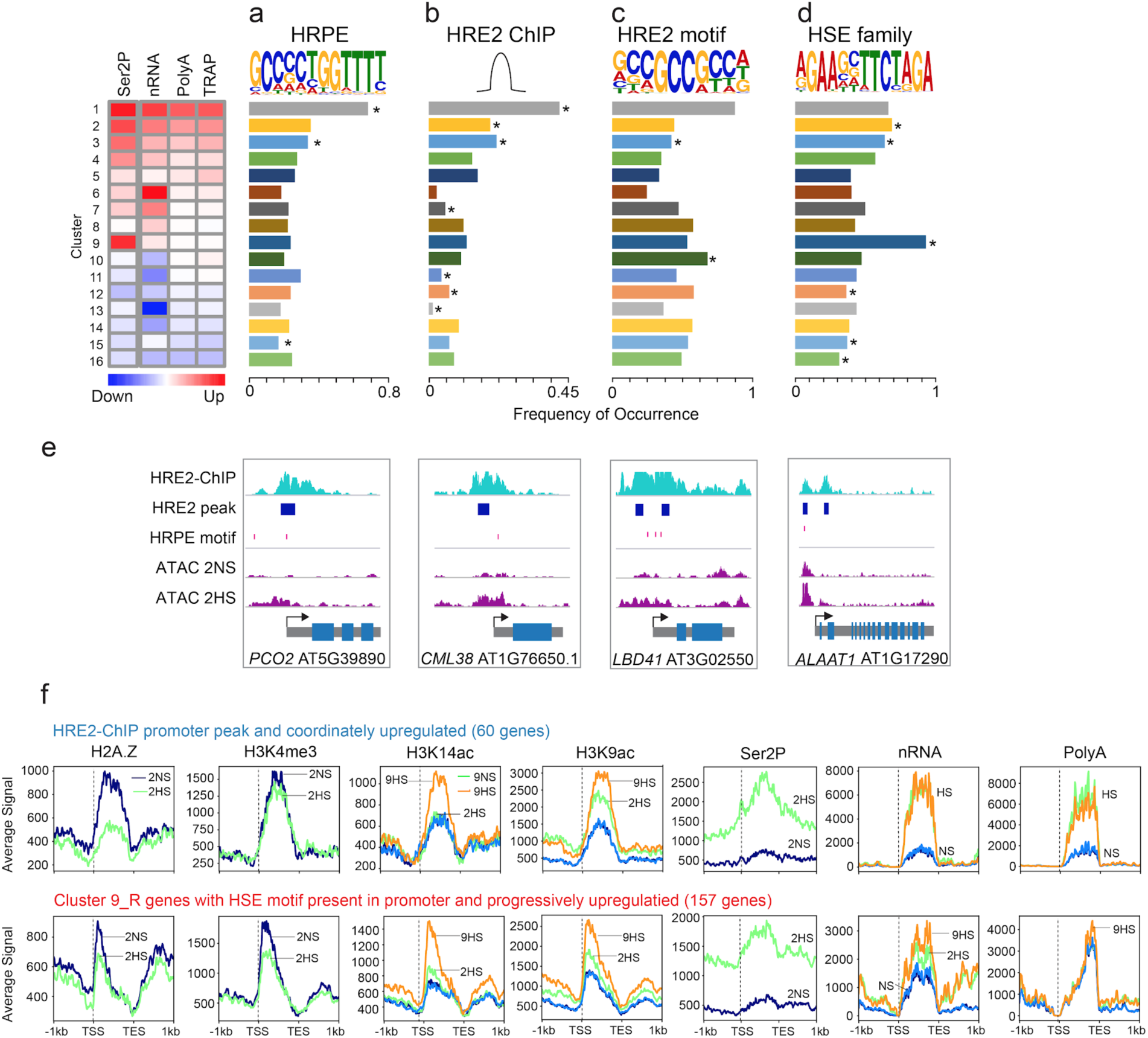
Regulation of hypoxic responses by two ERFVII and HSF TF families. **a-d** Motif and peak enrichment in the promoter regions (1 kb upstream to 500 bp downstream of the TSS) of genes in clusters identified in Figure 4c for **a**. Hypoxia Responsive Promoter Element (HRPE), **b**. HRE2-ChIP peaks, **c** HRE2-motif, and **d.** Heat Shock sequence Elements (HSE). Frequency of motif and peak occurrence was calculated as number of motifs per number genes in cluster. Clusters with significant enrichment were identified by two-tailed Fisher’s exact test p < 0.05. **e**. Genome browser view of select *HRGs* (*PLANT CYSTEINE OXIDASE 2, PCO2*; *CALMODULIN-LIKE 38, CML38*; *LOB-DOMAIN CONTAINING PROTEIN 41, LBD41*; *ALANINE AMINO TRANSFERASE 1, AAT1*). The locations of HRE2-ChIP peaks and HRPEs are depicted as a horizontal blue box and vertical red line, respectively. **f**. Average signal of chromatin and RNA readouts for **h,** HRE2 bound genes with coordinate upregulation and **i,** genes with progressive upregulation (cluster 9) containing promoter HSEs. Average signal is plotted from 1 kb upstream of the transcription start site (TSS) to 1 kb downstream of the transcript end site (TES).

The ERFVII HRE2 is encoded by an *HRG*. To gain insight into the role of HRE2, ChIP-seq was performed on 2HS seedlings overexpressing a stabilized version of this protein. Over 75% of the ∼1800 peaks identified mapped within 1 kb upstream of TSSs, with clear enrichment of binding within 500 bp upstream of the TSS (Supplemental Figure 10; Supplemental Table 2a), as evident in the promoters of *ADH1* and *PCO2* (Figure 5a,b; Supplemental Figure 10c,d). The binding of HRE2 to promoter regions was highest for cluster 1, but also significantly enriched for genes of clusters 2 and 3 of the 2HS analysis (Figure 7b). Overall, HRE2 peaks were present in 28% of the 213 coordinately up-DRGs (Supplemental Table 2).

We anticipated that HRE2 might be associated with the HRPE, known to be transactivated by RAP2.12, 2.2 and 2.3 (Gasch et al., 2015; Bui et al., 2015). But *de novo* motif discovery using the HRE2 peak regions yielded an overrepresented multimeric 5’-GCC-3’ element (p-value <1e^-351^) (Figure 7c; Supplemental Table 5) that resembles the GCC-rich EBP box bound RAP2.12 in the *in vitro* DNA affinity purification and sequencing (DAP-seq) assay (O’Malley et al., 2016)) and by RAP2.3 based on ChIP-qPCR and electrophoretic mobility shift assays (Büttner and Singh, 1997; Gibbs et al., 2014). The HRE2/GCC motif was enriched in cluster 1 but also was more prevalent than the HRPE in other clusters (Figure 7c). Our data strongly suggest that HRE2 binds promoter regions containing a GCC-rich EBP-like box. Yet, in over 50% of the 213 coordinately up-DRGs an HRPE is also present (Supplemental Table 2). Collectively, these observations support the conclusion that many of the coordinately up-DRGs are transcriptionally activated by one or more ERFVII.

### Hypoxic-stress progressively activates genes in heat and oxidative stress networks

The regulatory variation of oxidative stress and heat response genes displayed by clusters 2-3 and 9 genes at 2HS (Figure 4b) was of interest. Anoxia upregulates heat-stress associated genes (Banti et al., 2010; Loreti et al., 2005) and both the onset of hypoxia and reoxygenation trigger a reactive oxygen species (ROS) burst associated with signal transduction (Chang et al., 2012). Elevation of RNAPII-Ser2P without a commensurate increase in polyA RNA was characteristic of *HSP*s, *RESPIRATORY BURST OXIDASE D* and transcriptional co/activators associated with heat and oxidative stress (i.e., *MULTIPROTEIN BRIDGING FACTOR 1C* (*AtMBP1C*), *DEHYDRATION-RESPONSIVE 2A* (*DREB2A*), *ZINC FINGER PROTEIN 12* (*ZAT12*)). We also noted that several *HEAT SHOCK FACTOR* transcriptional activators were elevated in TRAP RNA at 2HS (*HSFA2, HSFA4A, HSFA7A*). We considered that the progressive upregulation of gene activity seen for cluster 9 genes may be mediated by HSFs and distinguished by specific chromatin features. Indeed, motif-enrichment analysis confirmed that HSF *cis*-elements (HSEs) were significantly enriched in promoters of cluster 9 (85% of genes) and to a lesser extent in cluster 2 and 3 genes (Figure 7d). Moreover, progressively up-DRGs with a 5’ HSE motif were distinguishable from the continuously upregulated HRE2-bound DRGs in four ways (Figure 7f): (1) a 5’ bias in nucleosomes, (2) more 5’ localized H2A.Z with less dramatic eviction at 2HS, (3) reduced H3K4me3 and increased H3K14ac that preceded the rise in H3K9ac, and (4) a delay in the increase in polyA RNA until 9HS. Thus, the regulatory variations for progressively up-DRGs with promoter-localized HSEs are distinct from those of the coordinately up-DRGs bound by HRE2 under hypoxia.

### Nuclear regulated stress responsive genes are highly translated upon reaeration

The discovery of genes with RNAPII-Ser2P engagement accompanied with 5’ biased nRNA but little to no polyA accumulation at 2HS (cluster 6; Figure 4b) indicates that some genes may be poised for upregulation upon release from oxygen deprivation. By evaluating dynamics in chromatin and RNA upon reaeration, the *HRG*s, *RP*s, and heat-responsive genes again showed distinct patterns of regulation (Figure 8a,b; Supplemental Figures 12-13). The co-clustering of all genes differentially expressed under hypoxia (2HS/2NS) or reaeration (R/2HS) identified cluster 8_R as upregulated in nRNA at 2HS but not in polyA RNA until R (Figure 8a,b; Supplemental Figure 12a). These genes were enriched for GO categories associated with heat (4.55e^-31^), high light intensity (1.92e^-33^), protein folding (8.91e^-33^), and reactive oxygen species (9.03e^-25^). Notably, over 50% of cluster 8_R genes had an HSE. The pattern of these HSE-containing genes contrasted to that of rapid progression towards non-stress conditions by *HRG*s and *RP*s in all three RNA readouts, as shown by the genome browser views of five representative genes (Figure 8c). The expression pattern demonstrated by HSE-containing genes may be physiologically relevant, as reaeration is associated with a rapid restoration of protein synthesis and a ROS burst. Together, our data demonstrate functional nuclear segregation of a subset of newly synthesized RNAs from cytoplasmic localized ribosomes in response to oxygen availability.

**Figure 8.**
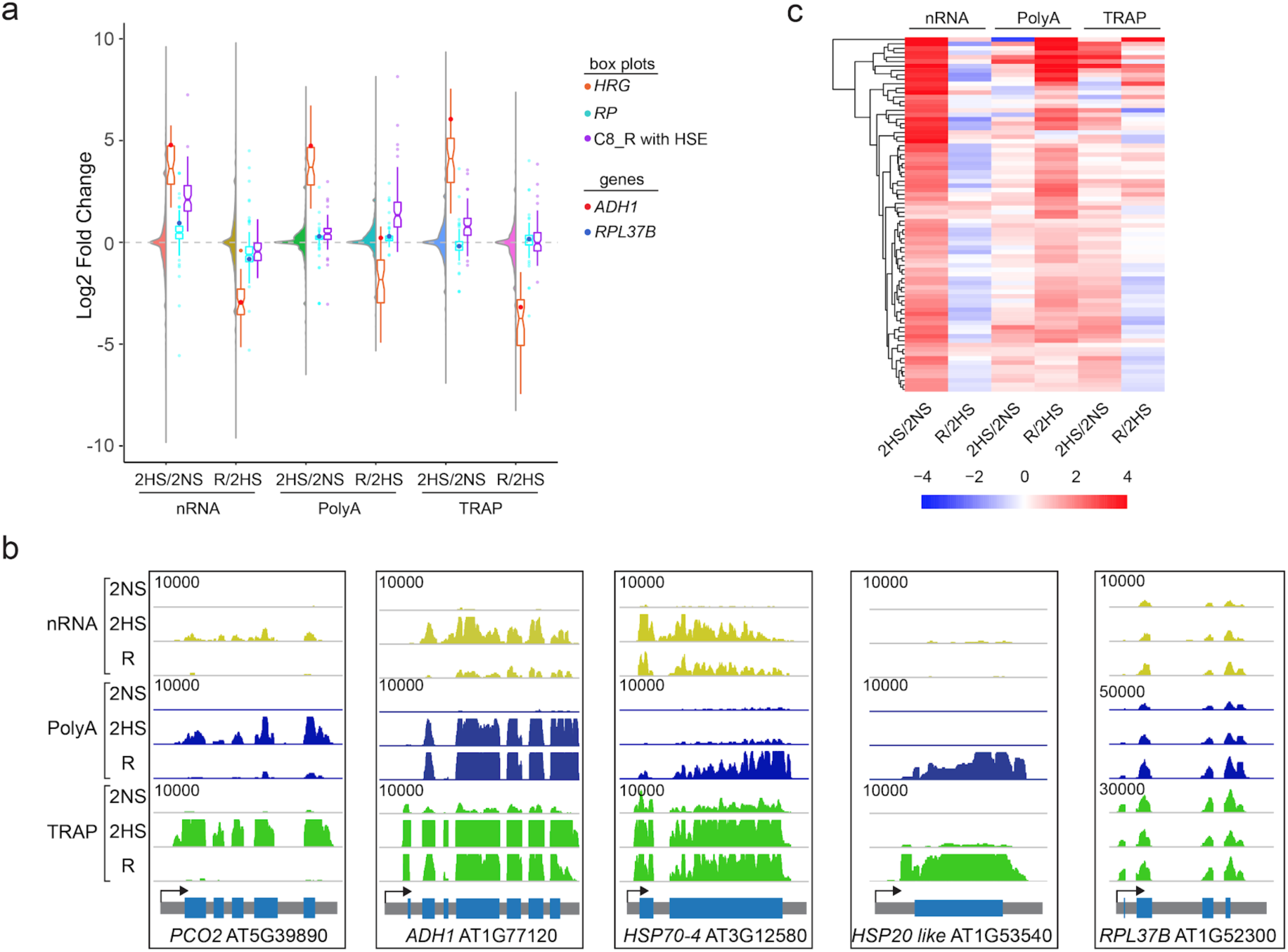
Reaeration promotes accumulation and translation of polyadenylated heat-responsive transcripts that are nucleus-enriched under hypoxia. **a** Combined violin and boxplot of stress-recovery dynamics of hypoxia-induced nuclear-enriched transcripts (2HS nRNA / 2NS nRNA, log_2_ FC > 3, n = 105). All genes shown as a violin; HRGs shown in orange, with *ADH1* as a red dot; RPs shown in blue with *RPL37B* as a blue dot; Heat Shock Element-containing cluster 8_R genes (C8_R, from Supplemental Figure 12) shown in purple. **b** Genome browser view of stress recovery dynamics of *HRGs* (*PCO2*, *ADH1*), heat responsive genes (*HSP70-4*, *HSP20 like*), and a representative *RP* (*RPL37B*). **c** Hierarchical clustering of Heat Shock Element (HSE)-containing cluster 8_R genes in response to hypoxia (2HS/2NS) and reoxygenation (R/2HS).

## Discussion

### Gene activity dynamics in response to hypoxic stress involves regulation of chromatin, transcription, and post-transcriptional processes

The dataset assembled of genome-wide measurements of chromatin state and intermediary steps of gene expression in seedlings exposed to sub-lethal hypoxic stress and reaeration identified time-dependent chromatin and transcript dynamics not appreciated by routine transcriptomics. Hypoxic stress modulated the (1) position and degree of open chromatin near the TSS, (2) modification of H3 lysines, (3) eviction of H2A.Z, (4) engagement of RNAP II-Ser2P, and (5) the abundance of nuclear, polyadenylated and ribosome-associated gene transcripts. The results illustrate clear distinctions between steady-state nuclear and polyA transcriptomes, as shown with similar methods for rice (Reynoso et al., 2017). The data also identified dynamics in chromatin accessibility and confirm translational dynamics in response to hypoxia and reaeration (Branco-Price et al., 2005, 2008a; Juntawong et al., 2014; Mustroph et al., 2009).

The monitoring of RNAPII-Ser2P engagement as a proxy for transcriptional elongation revealed that global levels of transcription were not dramatically dampened by 2HS (Figure 1c), although there was considerable regulation of individual genes (Figure 1c; Figure 4c). Remarkably, less than 10% of all DRGs were coordinately up- or downregulated from transcriptional elongation through translation at 2HS. Three patterns of discontinuous gene regulation were apparent: (1) progressive upregulation, characterized by high RNAPII-Ser2P engagement and delayed elevation of polyA RNA, as observed for heat and oxidative stress genes, (2) incomplete downregulation, characterized by maintained RNAPII-Ser2P engagement and increased nRNA abundance with limited change in polyA RNA, as observed for *RP*s; and (3) enhanced or reduced translational status, measured by comparison of TRAP RNA and polyA RNA abundance (Branco-Price et al., 2008a; Mustroph et al., 2009; Juntawong et al., 2014). We also identified variations in nuclear regulation (chromatin accessibility, histone modification, RNAPII engagement and nRNA abundance) that foreshadowed changes in polyA RNA abundance and translation. Finally, the evaluation of the three transcript populations following brief reaeration demonstrated a rapid rebound to homeostasis in the nRNA of the continuously up-DRGs, discontinuously up-DRGs and discontinuously down-DRGs. In sum, these data illuminate remarkable complexity in the transcriptional and post-transcriptional regulation of genes associated with stress-responses and growth.

### Coordinate upregulation of hypoxia-responsive genes is characterized by histone modification, histone variant eviction, and increased chromatin accessibility

Hypoxia had marked effects on the position, composition, and modifications of specific histones of nucleosomes. The genes that were coordinately upregulated from transcription through ribosome included the *HRG*s, critical to low-oxygen stress survival (Gibbs et al., 2011; Gasch et al., 2015; Mustroph et al., 2009, 2010; Giuntoli et al., 2014; Giuntoli and Perata, 2017). The strong and coordinate upregulation of *HRG*s was characterized by eviction of H2A.Z, a rise in gene body H3K9ac, promoter accessibility, RNAP-Ser2P engagement, nRNA, polyA and ribosome-associated mRNAs (Figure 9). The co-transcriptional placement of a positive charge on the N-terminus of H3K9 by histone acetyltransferases is a reliable signature of active transcription (Eberharter and Becker, 2002) and is common for stress-activated genes (Kim et al., 2008; Zhou et al., 2010; Kim et al., 2012). In animals, H3K9ac is promoted by H3K4me3 and stimulates the release of poised RNAPII complexes (Jonkers and Lis, 2015). H3K4me3 generally rose on *HRG*s in response to 2HS, but was not as strongly correlated with RNAPII-Ser2P engagement as H3K9ac (Figure 3b; Supplemental Figure 4d). The stress-activated transcription coincided with eviction of H2A.Z from nucleosomes that were broadly distributed across the bodies of *HRGs* at 2NS. This is consistent with prior reports that H2A.Z eviction from nucleosomes is a prerequisite for activation of temperature responsive genes (Kumar and Wigge, 2010; Sura et al., 2017; Cortijo et al., 2017; Dai et al., 2017; Torres and Deal, 2018).

**Figure 9.**
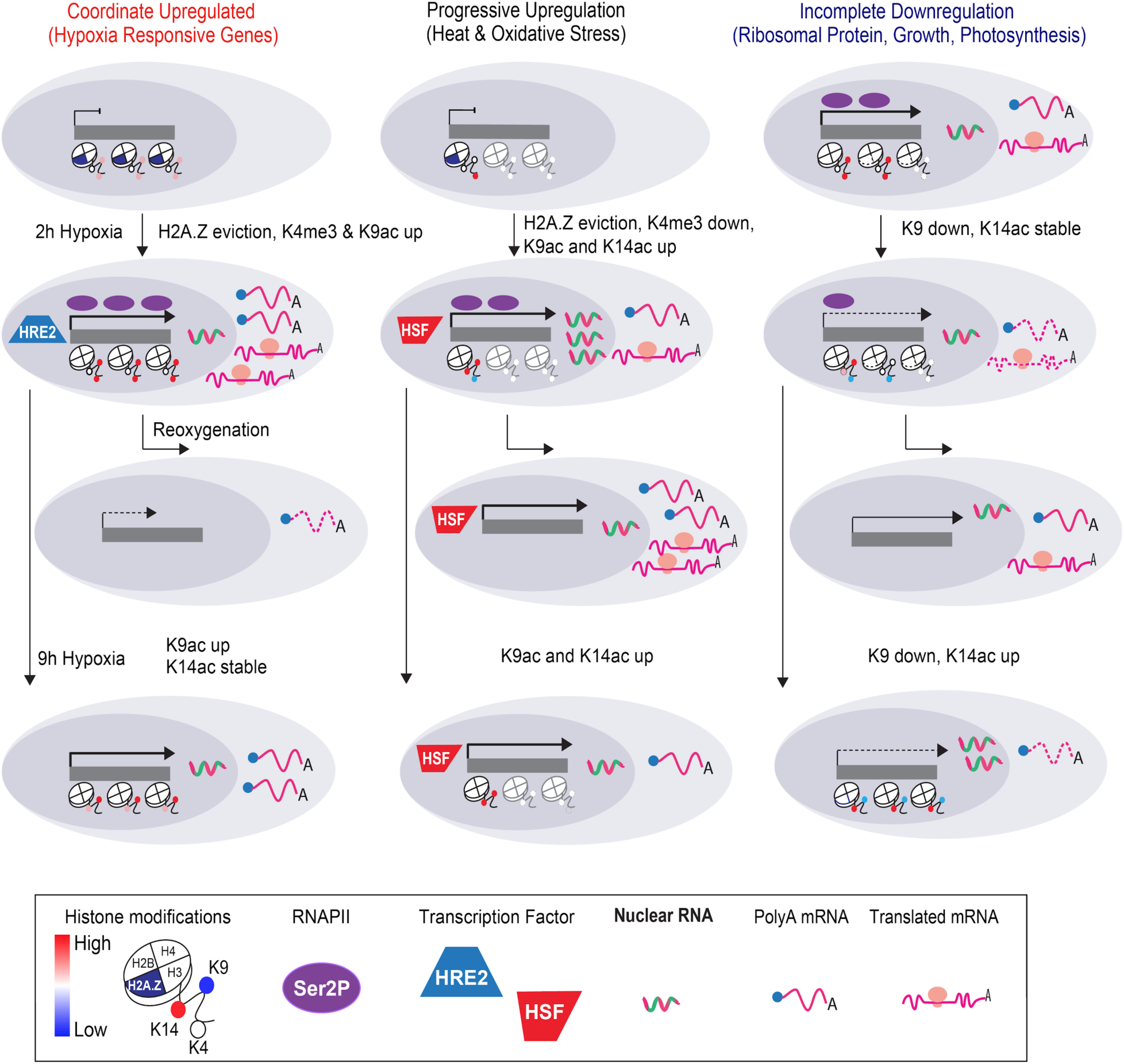
Three patterns of dominant nuclear regulation in response to hypoxic stress of two durations and reaeration. **(1)** Coordinate upregulation from chromatin to translation of hypoxia responsive genes characterized by hypoxia-triggered H2A.Z eviction, H3K4me3, and H3K9ac elevation. Over half of the genes with this pattern of regulation showed HRE2 binding by chromatin immunopurification. Transcript abundance and translation declines upon reoxygenation. (2) Progressive upregulation of genes associated with heat and oxidative stress responses characterized by a strong 5’ bias in histone marks, hypoxia-triggered H2A.Z eviction, H3K9ac elevation, and H3K4me3 reduction. Many are known targets of HSFs. These genes are lowly transcribed under normoxia (nonstress conditions), with evidence of transcriptional activation in response to brief hypoxia (2 h hypoxia) as shown by elevated RNAPII-Ser2P binding, nuclear (nRNA), and ribosome-associated transcripts (TRAP RNA), yet elevation in polyadenylated transcripts (polyA RNA) was limited until the stress was prolonged (9 h hypoxia). Transcript abundance increases upon reoxygenation. (3) Compartmentalized downregulation of many genes associated with growth and photosynthesis, including ribosomal protein genes characterized by maintained RNAPII-Ser2P and nRNA levels under the stress, reduced H3K9ac, and progressive acquisition of H3K14ac. Transcript abundance and translation is restored upon reoxygenation. Cartoons illustrate histone modifications as defined by the key. HRE2, Hypoxia-Responsive ERF 2; HSF, Heat Shock Factor transcriptional activator; Ser2P, RNAPII with the Ser2P modification.

H2A.Z deposition, eviction, and the impact on transcriptional regulation are complex and not fully understood. Deposition requires ACTIN-RELATED PROTEIN 6 (ARP6), a component of the SWR1 remodeling complex complex, and other factors in Arabidopsis (Asensi-Fabado et al., 2017; Kumar, 2018). Yet disruption of H2A.Z deposition in an *arp6* mutant has limited effect on regulation of *HRG*s under aerated growth conditions (Torres and Deal, 2018). Other chromatin remodeling proteins or specific TFs could be important. In response to small increases in temperature, displacement of H2A.Z was facilitated by HSF1A in the activation of the heat response network (Cortijo et al., 2017). H2A.Z eviction from *HRG*s could involve ERFVIIs. Indeed, the hypoxia-stabilized ERFVII RAP2.2 interacts with the SWI-SNF remodeler BRAHMA and contributes to H2A.Z eviction associated with increased chromatin accessibility (Efroni et al., 2013). Many of the HRGs displayed increased chromatin accessibility in their proximal promoter regions and gene bodies. Moreover, 69 of the >200 coordinately up-DRGs had one or more HRPE within 1 kb of their TSS and many were bound by HRE2 at 2HS (Figure 7e; Supplemental Table 2). RAP2.12 is stabilized and nuclear localized as external oxygen levels decline below 10 kPa (Kosmacz et al., 2015). The rise in nuclear levels of ERFVIIs under hypoxia may coordinate RNAPII initiation, facilitating H2A.Z eviction to promote productive elongation that is coupled with H3K9 acetylation. ERFVII interaction with the ATP-dependent chromatin remodeling machinery could also contribute to the expansion of regions of accessible chromatin to facilitate high levels of transcriptional activation. The uniform upregulation from transcription to translation of the up-DRGs with an HRPE and/or HRE2 bound promoter is reminiscent of the production of translationally competent mRNAs during nutrient starvation in yeast that is associated with specific transcription factors (Zid and O’Shea, 2014). We hypothesize that transcriptional activation by the ERFVIIs could promote the fidelity of co-transcriptional processing, export and translation of their targets under hypoxia.

### Genes associated with heat and oxidative stress are progressively upregulated by hypoxia

The stress distinctly activated genes associated with heat and oxidative stress. The progressive upregulation of cluster 9 genes after 2HS (Figure 4c; Supplemental Figure 7) became more evident when their activity was compared at 2HS and 9HS using two chromatin and two RNA readouts (Figure 7g). A significant proportion of these genes had HSEs within their promoters (Figure 4c, Figure 7d, Supplemental Figures 6 and 7). Their pattern of regulatory variation included a 5’ bias in histone marks and moderate to very high RNAPII-Ser2P engagement at 2HS accompanied by elevated nRNA and TRAP RNA, but limited full-length polyA RNA (Figure 8).

Previously, heat stress-responsive genes were recognized as highly induced in Arabidopsis seedlings directly transferred to anoxia (Loreti et al., 2005; Banti et al., 2010). Remarkably, a brief pre-treatment with heat stress increased the resilience to anoxia of wild-type seedlings but not loss-of-function *hsfa2* or *hsf1a hsf1b* mutant seedlings (Banti et al., 2008). This is enigmatic because high temperatures do not typically precede flooding in natural environments, but the availability of HSPs could reduce cellular damage under extreme stress conditions such as anoxia. Indeed, *HSPs* were plentiful in cluster 9 after 2HS. Limited synthesis of ATP-dependent chaperones after brief hypoxia by nuclear sequestration of their transcripts could minimize demands on limited energy reserves. However, as the stress is prolonged or upon reaeration, the rapid synthesis of chaperones and proteins that provide protection from ROS could aid survival. As an example, the upregulation of *RESPIRATORY BURST OXIDASE D* (*RBOHD*) contributes to survival of hypoxia, submergence and post-submergence recovery (Baxter-Burrell et al., 2002; Pucciariello et al., 2012; Gonzali et al., 2015; Yeung et al., 2018).

The nucleosome dynamics of cluster 2 and 9 genes after 2HS included similar modifications with notable exceptions (Figure 4c,d; Supplemental Figures 7 and 11). Genes of both clusters underwent a loss of H2A.Z near the TSS of the gene body in conjunction with increased H3K9ac, but cluster 9 genes showed a loss of H3K4me3 proximal to their TSS. This was not evident for cluster 2 or other coordinately up-DRGs. The decrease in H3K4me3 may be an indication of non-productive RNAPII-Ser2P engagement, as this modification is associated with RNAPII elongation (Kwak and Lis, 2013; Ding et al., 2012). Intriguingly, there was significant enrichment of HSEs in promoters of both rapidly and progressively upregulated genes (Figure 7d).

We predict that recruitment of specific TFs together with chromatin modifying enzymes may influence histone or RNAPII CTD modifications that influence the rate of RNAPII elongation, termination, or post-transcriptional processing, thereby tuning the temporal production of stress-induced transcripts. The progressive upregulation of heat and oxidative stress networks could be mediated by the upregulation of TF genes encoding HSFs (i.e., HSFB2B) and ZINC FINGER PROTEIN 10 (ZAT10), and others during the early hours of the stress. Notably, the level of polyA mRNA accumulation and translation of these mRNA was enhanced upon reaeration (Figure 8a). Our datasets provide an opportunity for further meta-analyses of heat and oxidative stress gene regulatory networks.

### Genes associated with major cellular processes continue to be transcribed during hypoxia

Intriguing patterns of transcriptional and RNA regulation of genes associated with processes associate with growth became evident from our multi-tier evaluation (Figure 8). Our prior comparisons of the transcriptome and translatome determined that hypoxia limits the investment of energy into the production of abundant cellular machinery such as ribosomes. mRNAs encoding *RPs* and other abundant proteins were seen to be maintained but poorly translated under hypoxic stress (Branco-Price et al., 2008a, 2005; Juntawong et al., 2014; Sorenson and Bailey-Serres, 2014). The stress-limited translation of *RP*s mRNAs is associated with their sequestration in UBP1C-associated macromolecular complexes (Sorenson and Bailey-Serres, 2014). Upon reaeration, *RP* mRNAs rapidly re-associate with polysomes (Branco-Price et al., 2008a; Juntawong et al., 2014; Sorenson and Bailey-Serres, 2014). This is the first study to consider at the global scale if *RP* or other translationally limited mRNAs are synthesized during hypoxia, although nuclear run-on transcription assays determined that housekeeping gene transcriptions continues in roots of fully submerged maize seedlings (Fennoy et al., 1998). Our RNAP-Ser2P and nRNA data confirmed that genes encoding RPs, metabolic enzymes and some photosynthetic apparati are synthesized and retained as partially or completely processed transcripts in the nucleus during hypoxia (Figure 6), a situation readily reversed by re-aeration (Figure 8a). These genes showed progressive increases in H3K14ac but not H3K9ac between 2HS and 9HS. The limited increase in their polyA RNAs could be due to limited cleavage and polyadenylation or increased turnover. Similar uncoupling of the H3K9ac and H3K14ac marks associated with active transcription but limited polyA RNA accumulation was reported for promoters deemed inactive but primed for future activation in pluripotent embryonic stem cells of mice (Karmodiya et al., 2012). The incomplete downregulation of housekeeping genes may poise cells for the observed rapid recovery upon reaeration.

Our genome-scale analysis demonstrate retention of transcripts within the nucleus during hypoxia, consistent with visualization studies of specific mRNAs in Arabidopsis and lupine (Niedojadło et al., 2016). mRNAs associated with the cell cycle are reportedly enriched in the nucleus under control conditions in Arabidopsis (Yang et al., 2017). This was confirmed in our comparison of the nRNA and polyA RNA transcriptomes under control and hypoxia stress (Figure 5). These data support a scenario wherein the nuclear compartmentalization of abundant transcripts contributes to energy conservation by limiting translation, similar to the cytoplasmic sequestration associated with the RNA binding protein UBP1C. Nuclear retention and cytoplasmic sequestration could provide two reservoirs of transcripts that can be deployed upon reaeration, enabling rapid restoration of cellular processes. Indeed, our data provide evidence that nuclear retained transcripts increase in translation status upon reaeration (Figure 8a).

Thus, the comparative monitoring of nRNA, polyA and TRAP RNA populations strongly support the conclusion that nuclear export and subsequent accumulation and translation of mRNAs is dynamically regulated by stress in plants. Factors such as chromatin accessibility, RNAPII elongation and termination and histone modification (i.e., H3K14ac) could contribute to dynamic in nuclear transcript abundance. Circumstantial evidence of altered co-transcriptional processing includes hypoxia-promoted intron retention (Juntawong et al., 2014) and alternative polyadenylation site selection (de Lorenzo et al., 2017). Although there is no clear evidence of a general disruption of splicing or polyadenylation during hypoxia (Koroleva et al., 2009; Juntawong et al., 2014; de Lorenzo et al., 2017), these processes could be altered by changes in chromatin, RNAPII elongation or association with certain proteins The RNA helicase eIF4AIII (Koroleva et al., 2009) and several RNA binding proteins (UBP1A/B/C and CML38) form macromolecular complexes present in the nucleus but primarily cytoplasmic under hypoxia (Sorenson and Bailey-Serres, 2014; Lokdarshi et al., 2016). Further experimentation is required to better understand the controlled dynamics in transcript synthesis, processing and export during hypoxia stress.

### An ERFVII hierarchy drives transcriptional activation in response to hypoxia

This study included the first genome-wide analysis of the *cis*-element targets of an ERFVII. ChIP-seq of the hypoxia-induced ERFVII HRE2 identified a multimeric 5’-GCC-3’ *cis*-element that is distinct from the HRPE that is enriched in *HRG* promoters and activated by ERFVIIs (Gasch et al., 2015). This is consistent with the observation that neither HRE1 nor HRE2 interacted with a 33 bp promoter fragment containing the HRPE in a Yeast-1 hybrid assay or provided significant transactivation via the HRPE in protoplasts (Gasch et al., 2015). The HRE2 binding motif strongly resembled the double GCC motif of the EBP box that was first defined for an ERF of *Nicotiana tabacum* (Sato et al., 1996). Previously, RAP2.3 was shown to bind the *ACID INSENSITIVE 5* (*ABI5*) promoter in a region with two EBP boxes; transactivation of the *ABI5* promoter in protoplasts by all three constitutively expressed ERFVIIs (RAP2.2/2.3/2.12) was dependent on the EBP box (Gibbs et al., 2014).

Collectively, our observations support earlier conclusions that members of the two ERFVII subgroups are not functionally redundant (Gasch et al., 2015; Gibbs et al., 2014). Yet, the situation remains somewhat enigmatic. The HRPE contains a sequence resembling a double 5’-GCC-3’. Moreover, promoter regions with high scores for the HRPE and HRE2 motif coincided in 40 of the 213 coordinately up-DRGs, of which 13 were bound by HRE2 (Supplemental Table 1) and the Arabidopsis *ADH1* promoter region that is sufficient for hypoxic upregulation contains both of these motifs (Gasch et al., 2015). The current model is that the low-oxygen-mediated stabilization of the constitutively-expressed RAP-type ERFVIIs is responsible for rapid transactivation of *HRG*s that include *HRE2* (van Dongen and Licausi, 2015; Voesenek and Bailey-Serres, 2013), which is synthesized and stabilized under hypoxia (Gibbs et al., 2011). We propose that HRE2 production and binding to promoters via the multimeric GCC element (HRE2 motif) is important for continued upregulation of *HRG*s, as an *hre1 hre2* double mutant fails to maintain the upregulation of these genes after 4 h of anoxia (Licausi et al., 2010). It may also enable cells to circumvent the inhibition of RAP2.12 activity by its interaction with the HYPOXIA RESPONSE ATTENUATOR (HRA1) (Giuntoli et al., 2014). Further analyses are required to comprehend the independent or overlapping roles of the RAP-type and HRE ERFVIIs in the temporal activation of transcription during hypoxia.

Mammals regulate transcriptional activity in response to hypoxia in a manner that is remarkably analogous to Arabidopsis (Watson et al., 2010). Hypoxia-responsive genes are activated by the two-subunit Hypoxia Inducible Factor (HIF) TF complex, that includes the oxygen-destabilized HIF1A subunit (LaGory and Giaccia, 2016). Similar to the three RAP-ERFVIIs of Arabidopsis, HIF1A is a constitutively synthesized protein that is destabilized in the presence of oxygen. A group of prolylhydroxlases catalyze the oxygen-dependent hydroxylation of specific residues of HIF1A that triggers its ubiquitylation and protoeasome-mediated turnover. In animals, the chromatin landscape is additionally regulated through chromatin accessibility (Buck et al., 2014) and the post-translational modification of histones (Chervona and Costa, 2012) in an oxygen-sensitive manner. HIF signaling itself directly facilitates histone modification of target genes through interactions with chromatin modifying enzymes (Watson et al., 2010), several histone modifying enzymes are directly targeted by HIF transcriptional activation, and HIF1A expression itself is regulated by histone acetylation (Kim et al., 2007). Moreover, specific histone modifying enzymes function as direct oxygen sensors through the mediation of their activity based on cellular oxygen levels (Batie et al., 2019; Chakraborty et al., 2019), confirming the integration of oxygen sensing with epigenetic regulation. The low-oxygen stabilized transcriptional regulators and the coordination of their function with chromatin modifications in Arabidopsis is evidence of convergence in gene activity regulation in response to hypoxia in plants and animals. This may extend beyond transcriptional regulation as cytoplasmic sequestration and selective translation of mRNAs is also observed in plants and metazoa (Blais et al., 2004; Young et al., 2008; Branco-Price et al., 2008a; Juntawong et al., 2014). It remains to be explored if hypoxia influences the synthesis and accumulation of nascent transcripts associated with growth in animals, as demonstrated in this multi-omic analysis of Arabidopsis.

### Conclusions

This multi-level investigation into gene regulatory processes revealed unexpectedly complex regulation of gene activity within the nucleus and cytoplasm that fine tune cellular and physiological acclimation to transient hypoxia, providing a model for response to an acute stress. A key finding is that there are rapid changes in the epigenome in response to short-term hypoxic stress that continue as the stress becomes more severe. Coordinate up-regulation from chromatin accessibility to translation was observed for over 200 hypoxia-responsive genes. Extreme stress-responsive and growth-associated genes showed more discontinuous upregulation of nascent transcript production, export to the cytoplasm, and recruitment to ribosomes. Each of these broadly defined gene cohorts showed a distinct pattern of epigenetic, nuclear and cytoplasmic transcript regulation upon re-aeration. Future investigations can consider roles of specific TF and their interactions with RNAPII, transcriptional coactivators, the chromatin landscape, and the post-transcriptional apparatus in directing the nuanced nuclear regulation uncovered in this study.

## Materials and Methods

### Genetic Material

The following genotypes were used for genome wide experiments: Col-0, Histone modification ChIP-seq, RNAP II ChIP-seq; *pHTA11:HTA11-3xFLAG (Cortijo et al., 2017)*, H2A.Z ChIP-seq; *pUBQ10:NTF/pACT2:BirA (Sullivan et al., 2014)*, ATAC-seq, nRNA-seq, polyA mRNA-seq; *p35S:HF-RPL18* (Zanetti et al., 2005), TRAP-seq, mRNA-seq; and HRE ChIP-seq *p35S:C2A-HRE2-HA* (Gibbs et al., 2011).

### Plant Growth and Treatment

Arabidopsis seeds were sterilized by incubation in 70% EtOH for 5 min, followed by incubation in 20% (v/v) bleach, 0.01% (v/v) TWEEN-20, followed by three washes in ddH_2_O for 5 min, in triplicate. Sterilized seeds were placed onto 1x MS media (1.0x Murishige Skoog (MS) salts, 0.4% (w/v) Phytagel (Sigma-Aldrich), and 1% (w/v) Suc, pH 5.7) in 9 cm^2^ Petri dishes and stratified by incubation at 4°C for 3 d in complete darkness. Following stratification, plates were placed vertically into a growth chamber (Percival) with 16 h light / 8 h dark cycle at ∼120 μmol photons·s^-1^·m-^2^, at 23 °C for 7 d. For hypoxic stress, seedlings were removed from the growth chamber at Zeitgeber time (ZT) 16 and were subjected to hypoxic stress by bubbling argon gas into sealed chambers in complete darkness for 2 or 9 h at 24 °C. The chamber set-up was as described by (Branco-Price et al., 2008a). Oxygen partial pressure in the chamber was measured with the NeoFox Sport O_2_ sensor and probe (Ocean Optics). Hypoxia ([O_2_] < 2%) was attained within 15 min of initiation of the stress. Control samples were placed in an identical chamber that was open to ambient air under the same light and temperature conditions. For reaeration, seedlings were identically subjected to hypoxic stress for 2 h and subsequently placed into the control chamber for 1 h (nRNA and polyA) or 2 h (TRAP) following the stress. Tissue was rapidly harvested into liquid N_2_ and stored at −80 °C.

### Chromatin and Nuclei Preparation

#### Detailed methods are provided in the supplement

ChIP was performed according to (Deal and Henikoff, 2010) with minor modifications provided in the supplemental methods. INTACT was performed according to (Deal and Henikoff, 2010) with modifications provided in the supplemental methods. ATAC-seq was performed on fifty-thousand INTACT-purified nuclei from root tissue according to (Buenrostro et al., 2013). Nuclear RNA extraction of INTACT-purified nuclei was performed according to (Reynoso et al., 2017). Additional details are provided in Supplemental File 1.

### Nuclear, PolyA and TRAP-RNA Isolation and Library Construction

Total RNA was extracted from 50 mg frozen pulverized tissue using Trizol (Life Sciences), treated with DNaseI, purified using AMPure XP beads, and eluted in 50 µL H_2_O and polyA selected. PolyA RNA selection was performed using Oligo (dT)_25_ Dynabeads by two rounds of magnetic capture (Townsley et al., 2015). TRAP (Translating Ribosome Affinity Purification) of mRNA-ribosome complexes was performed following the procedure of (Mustroph et al., 2009). TRAP RNA extraction (Townsley et al., 2015) was performed with minor modifications and then polyA RNA was selected as described by (Townsley et al., 2015). RNA-seq library preparation for nRNA, polyA, and TRAP RNA was performed according to (Kumar et al., 2012). Additional details are provided in Supplemental File 1.

### Bioinformatic Analyses

For all high throughput outputs, short reads were trimmed using FASTX-toolkit to remove barcodes and filter short and low quality reads prior to alignment to the TAIR10 genome with Bowtie2/Tophat2. Read quality reports were generated using fastqc. For all datasets excluding ATAC-seq and HRE2 ChIP-seq, counting of aligned reads was performed on entire transcripts (Lawrence et al., 2013) using the latest Araport 11 annotation (201606) and reads per kilobase of transcript per million mapped reads (RPKM) values were calculated. Differentially expressed genes were determined by edgeR, |FC| > 1, FDR < 0.01. *Data visualization and file generation*: Bigwig files for Integrated Genome Viewer (IGV) visualization were generated from aligned bam files and normalized by RPKM values using the deepTools command bamCoverage with the - normalizeUsingRPKM specification. Within IGV, all chromatin based outputs were normalized to the same scale and all RNA outputs were normalized to a separate scale. *Peak calling:* For ChIP-seq and ATAC-seq datasets, peak calling was performed using independent replicates in combination as input for HOMER using the findPeaks command (Heinz et al., 2010). For downstream analyses, peaks identified from each time point comparison were combined and counting was performed on individual replicates. Peaks were annotated using the HOMER command annotatePeaks.pl. Differential regulation of peaks was performed using edgeR. *Motif discovery:* HRE2 peaks identified by HOMER command findPeaks were used as an input for the HOMER command findMotifsGenome.pl. The matrix for the top HRE2 motif (p: 1e-69) was used as input the MEME command ceqlogo to generate the motif logo. *DAP-seq motif analysis:* Sequences of promoter regions spanning 1 kb upstream to 500 bp downstream of the TSS were extracted for genes within each cluster. The promoter sequences were compared against each TF, per TF family, using motif matrices identified by (O’Malley et al., 2016). The number of significant motifs identified within the promoters of each cluster were quantitated and normalized to the number of genes within each cluster. *t-distributed stochastic neighbor embedding (t-SNE)*: t-SNE analysis was performed according to (Cao et al., 2017). Briefly, a principal component analysis was performed using RPKM values for each replicated dataset and timepoint. t-SNE was then performed on PCs 1-10 and the results were plotted using ggplot2.

*Wilcoxon signed rank sum test:* Wilcoxon signed rank sum test was performed by comparing the RPKM values of one genomic output for each gene within a cluster to the genes of another cluster. Each cluster was compared individually to all other clusters to determine significance, p < 0.05. *Fisher’s exact test:* Fisher’s exact test was performed by comparing the number of motifs / peaks associated with genes within a cluster to the total number of motifs / peaks identified for all DRGs.

### Bioinformatic Tools

Analyses were performed with Bioconductor R packages particularly the Next Generation Sequencing analysis software of systemPipeR (Girke, 2014). Programs used within Bioconductor included: BiocParallel (Morgan et al., 2014), BatchJobs (Bernd Bischl et al., 2015), GenomicFeatures (Lawrence et al., 2013), GenomicRanges (Lawrence et al., 2013), edgeR (McCarthy et al., 2012), gplots (Warnes et al., 2009), ggplot2 (Wickham, 2009), RColorBrewer (Neuwirth, 2011), Dplyr (Wickham and Francois, 2015), biomaRt (Durinck and Huber), ChIPseeker (Yu et al., 2015), rtracklayer (Lawrence et al., 2009), Rtsne (Krijthe, 2015), Pheatmap (Kolde, 2012), e1071 (Meyer et al., 2015), cluster (Maechler, 2018), ShortRead (Morgan et al., 2009), clusterProfiler (Yu et al., 2012), and Rsamtools (Morgan et al., 2016). The Publically available UNIX/Python/Perl packages used included Tophat (Kim et al., 2013), fastx_toolkit (Gordon and Hannon, 2010), Fastqc (Andrews and Others, 2010), MEME (Bailey et al., 2009), HOMER (Heinz et al., 2010), deepTools (Ramírez et al., 2016), circos (Krzywinski et al., 2009), Bedtools (Quinlan and Hall, 2010), samtools (Li, 2011).

### Statistical analysis

Statistical analysis of cluster motif enrichment was performed by two-tailed Fisher’s exact test comparing motif frequency for individual genes of each cluster; p <0.05. Statistical significance of cluster values was performed for each genomic output using Wilcoxon signed rank test for values of all genes within each cluster. p < 0.05.

## Accession Numbers

The Arabidopsis genes examined in this study include: ADH1, AT1G77120; PCO2, AT5G39890; HSP70-4, AT3G12580; RPL37B, AT1G52300; LHY1, AT1G01060; CCA1, AT2G46830; PRR9, AT2G46790; PRR7, AT5G02810; PRR5, AT5G24470; RV8, AT3G09600; LUX, AT3G46640; CIR1, AT5G37260; ELF3, AT2G25930; ELF4, AT2G40080; LWD1, AT1G12910; ZTL, AT5G57350; GI, AT1G22770; LNK1, AT5G64170; LNK2, AT3G54500; TOC1, AT5G61380; CML38, AT1G76650; LBD41, AT3G02550; ALAAT1, AT1G17290; HSP20 like AT1G53540;

## Supplemental Data

Supplemental Figure 1. Expanded overview of the multiscale chromatin and RNA gene regulatory analyses.

Supplemental Figure 2. Average signal plots and heat maps for the global distribution of reads of chromatin and RNA data.

Supplemental Figure 3. Multi-level evaluation of histone and gene regulation activity of the *HRG*s and *RP*s.

Supplemental Figure 4. Evidence of cooperative association between H2A.Z eviction and H3K9ac and the relation between Histone H3 modification and RNAPII engagement.

Supplemental Figure 5. Genome-wide comparison of nRNA to polyA RNA demonstrates nuclear and cytoplasmic enrichment of transcripts under two conditions.

Supplemental Figure 6. Distinctions in histones, RNAPII-Ser2P and RNA levels on genes that are co-regulated in response to hypoxic stress.

Supplemental Figure 7. Progressive regulation of variations in gene activity in response to hypoxia.

Supplemental Figure 8. Histone and gene activity comparisons of genes co-regulated after 9 h of hypoxic stress.

Supplemental Figure 9. Quantitative analysis of dynamics in histone modifications of co-regulated gene clusters after 2 h of hypoxic stress.

Supplemental Figure 10. Genome-wide binding dynamics of HRE2 determined by ChIP-seq.

Supplemental Figure 11. Genome browser view of normalized read coverage of histone, ATAC-seq, RNAPII-Ser2P, HRE2-chromatin immunopurification and RNA outputs for representative genes of cluster 9.

Supplemental Figure 12. Reoxygenation following hypoxia promotes global dynamics in chromatin accessibility and RNA populations.

Supplemental Figure 13. Regulation of variations in gene activity in response to hypoxia and reoxygenation.

Supplemental Table 1. Tabulation of ATAC and HRE2-ChIP peaks on chromatin

Supplemental Table 2. Survey of histones, RNAPII, HRE2 and three RNA sub-populations in response to two durations of hypoxic stress

Supplemental Table 3. nRNA Enrichment and Gene Ontology Analysis

Supplemental Table 4. Gene Ontology Analysis for two durations of hypoxic stress in

Supplemental Table 5. List of Arabidopsis Blacklisted Chromatin (ABC)

Supplemental File 1. Supplementary Methods

Supplemental File 2. BED file of Arabidopsis Blacklisted Chromatin (ABC)

## Supplemental Note

### *Arabidopsis* blacklisted chromatin

The variety of genome-wide outputs in this study recording dynamics at the level of chromatin and RNA has permitted the identification of Tn5 bias/genomic regions with high nonspecific background and/or high signal/read counts in Arabidopsis irrespective of experiment, similar to regions identified by the ENCODE project for other organisms (ENCODE Project Consortium, 2012). The identification and filtering of genomic blacklisted regions is especially important for calculation of genome wide Pearson correlations and the generation of signal tracks used for accurate browser views. The *<u>A</u>rabidopsis* <u>B</u>lacklisted <u>C</u>hromatin (ABC) regions (Supplemental Table 5, Supplemental File 2) can be used to remove these defined regions that create statistical artefacts of the nuclear genome prior to performing analyses of chromatin-based data.

## Acknowledgements

We thank Mauricio Reynoso, Roger Deal, Marko Bajic, Seung-Cho Lee, Maureen Hummel, Lauren Dedow, Sonja Winte, Jérémie Bazin, Thomas Girke, and Thomas Eulgem for helpful discussions. This work was supported by U.S. National Science Foundation (Grant Nos. IOS−1121626, IOS-1238243, MCB-1716913) and the U.S. Department of Agriculture, National Institute of Food and Agriculture - Agriculture and Food Research Initiative (Grant No. 2011−04015) to J.B.-S.

## Author contributions

T.L. and J.B.-S. designed research; T.L. performed research; T.L. and J.B.-S. analyzed data; and T.L. and J.B.-S. wrote the paper.

---

**Supplemental Figure 1.**
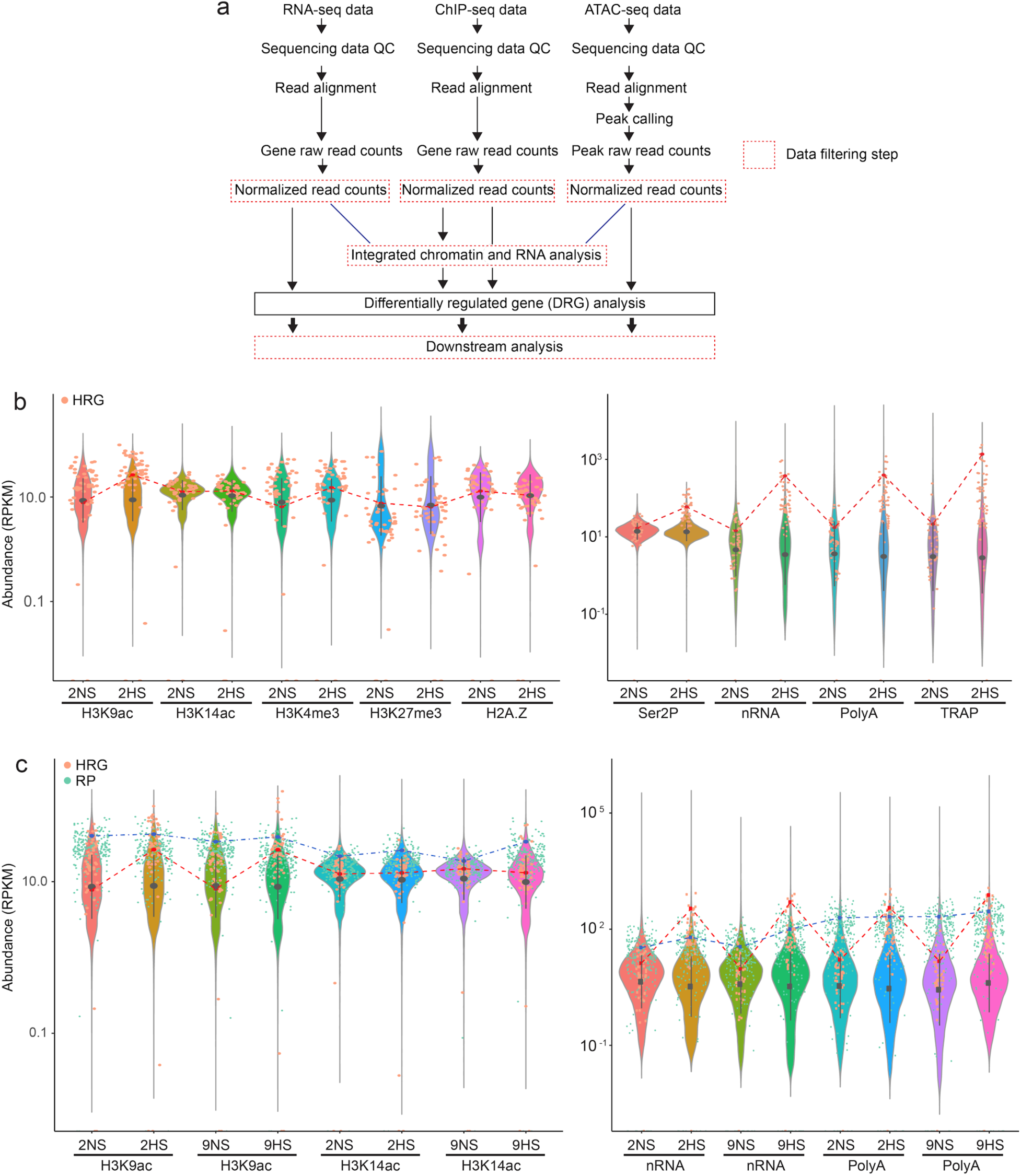
Expanded overview of the multiscale chromatin and RNA gene regulatory analyses. **a** Schematic of library types and data analysis performed on 7-day-old seedlings subjected to control (normoxic) (2NS, 9NS) or hypoxic stress (2HS, 9HS) as shown in Figure 1a. Control seedlings were compared to hypoxia stress seedlings independently within each chromatin- and RNA-based dataset. Low abundance reads were filtered (< 2 CPM). Differentially regulated genes were identified and utilized for downstream analyses. The chromatin and RNA datasets were also analyzed in combination. **b-c** Violin plots comparing distribution of genome-wide abundance (reads per kilobase per million reads [RPKM]) of histone and mRNA-related outputs for control (2NS, 9NS) and hypoxic (2HS, 9HS) treatments for all readouts assayed at 2HS **b**, and 9HS **c.** RPKM values for the 49 core hypoxia responsive upregulated genes (HRGs) (Mustroph et al., 2009) are plotted as orange dots. The HRG, *ALCOHOL DEHYDROGENASE 1* (*ADH1*) is tracked as a red line. RPKM values for cytosolic ribosomal proteins (RPs) are plotted in blue, with *RIBOSOMAL PROTEIN 37B* tracked between datasets (blue dashed line). Mean ± SD are depicted within violins as a dark gray dashed line.

**Supplemental Figure 2.**
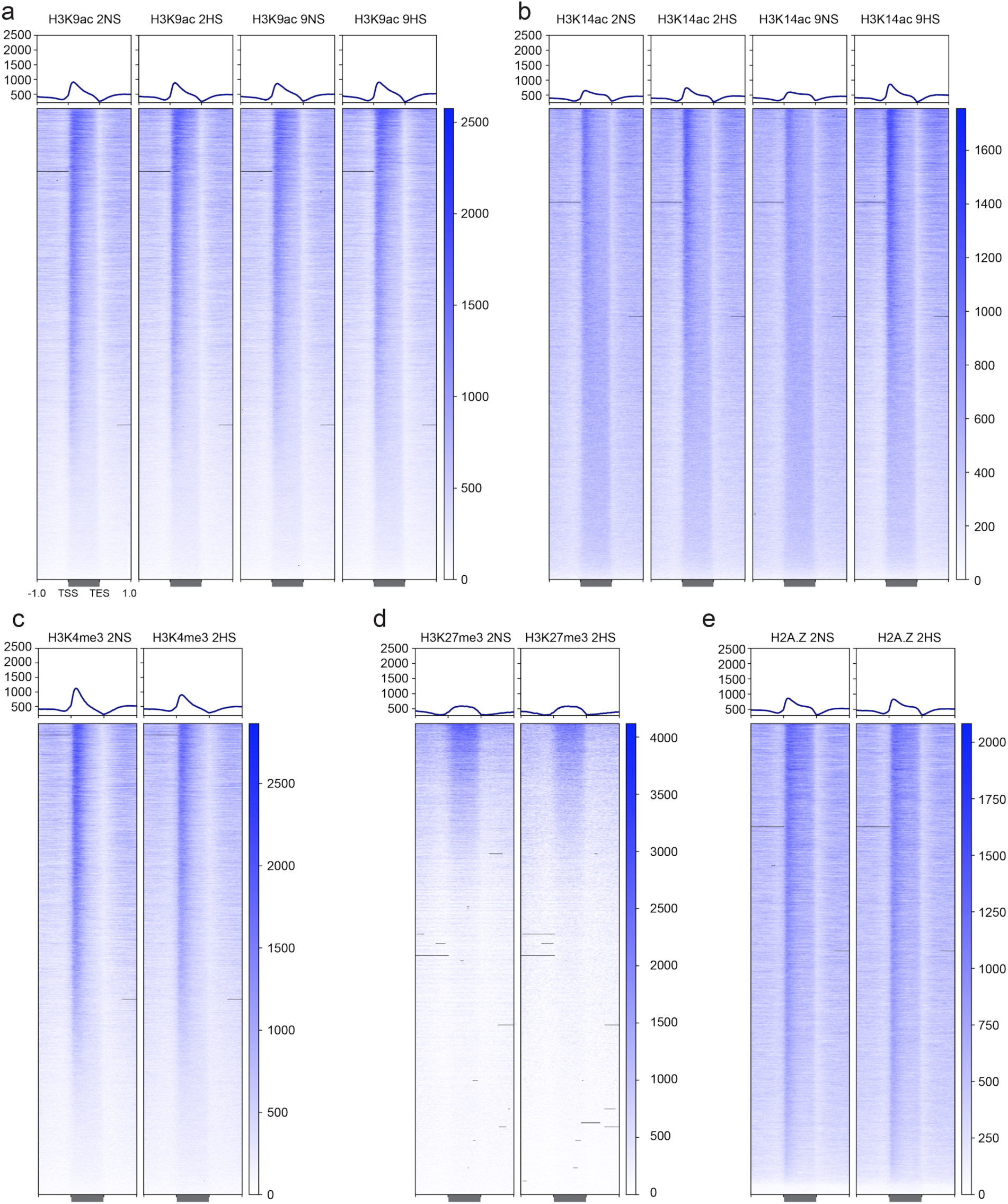
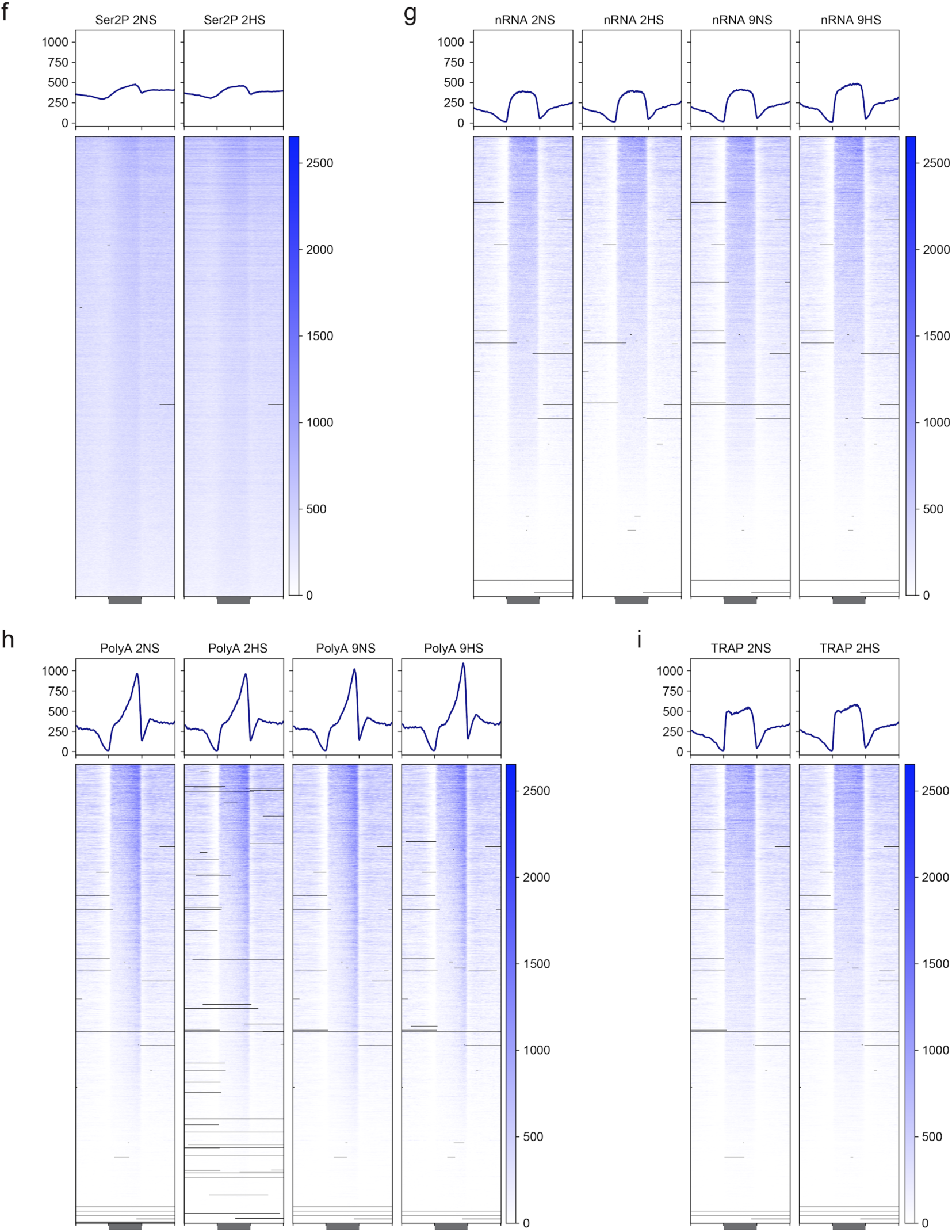
Average signal plots and heat maps for the global distribution of reads of chromatin and RNA data. **a-h** The genomic region from 1 kb upstream of the transcription start site (TSS) to 1 kb downstream of the transcription end site (TES) is depicted for each histone modification and variant, RNAPII-Ser2P (Ser2P), nuclear RNA (nRNA), polyadenylated RNA (polyA RNA) and ribosome-associated RNA (TRAP RNA) for nonstress (2NS, 9NS) and hypoxia (2HS, 9HS) stressed seedlings. Each heat map ranks genes from the highest signal (top) to the lowest (bottom). Heatmap intensity scale values differ between plots of different readouts and are shown. Blacklisted regions of the genome with disproportionately high reads are tabulated in Supplemental Table 5 and depicted as a black line.

**Supplemental Figure 3.**
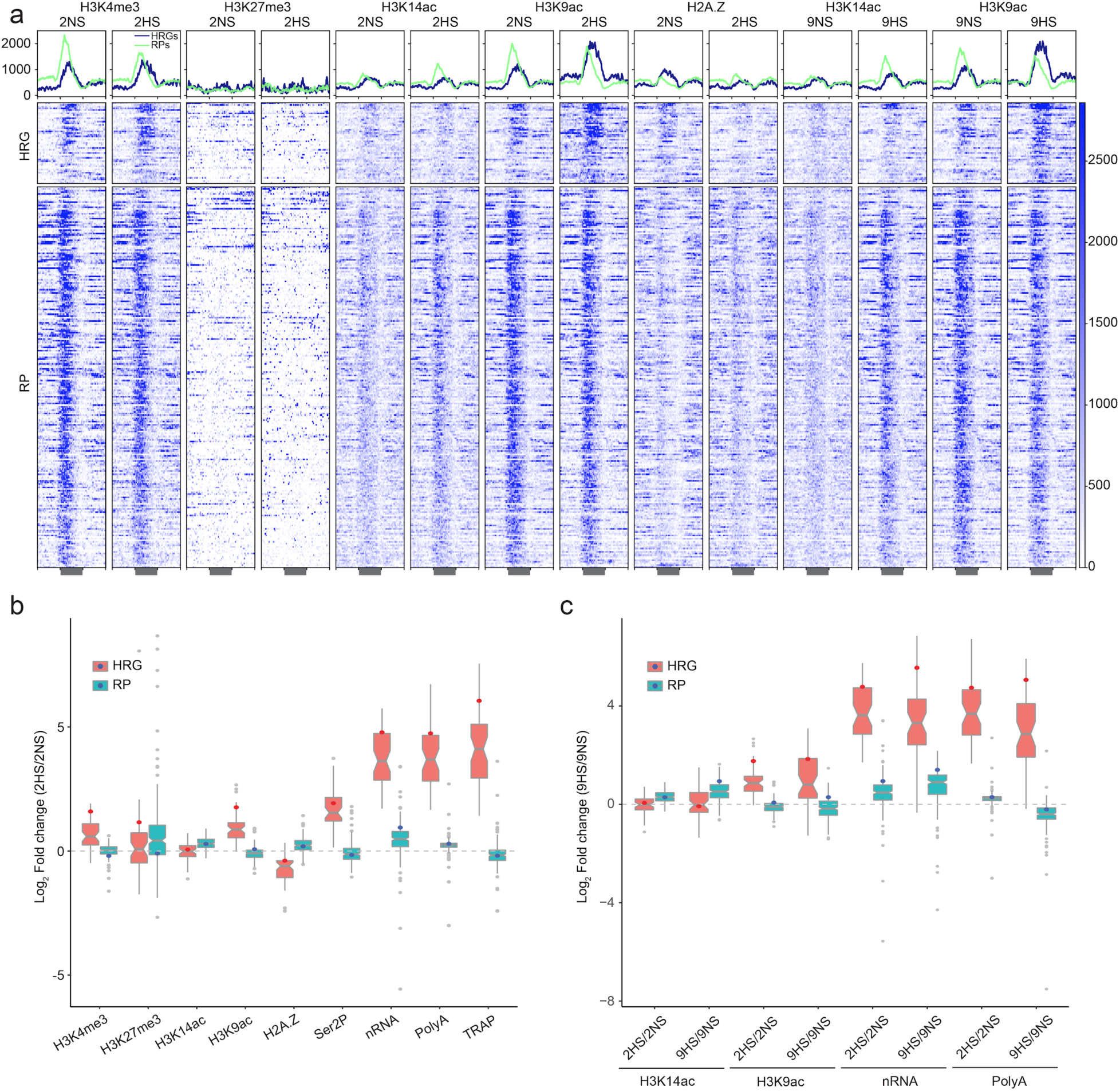
Multi-level evaluation of histone and gene regulation activity of the *HRG*s and *RP*s. **a** The average signal profile and read distribution heatmap for *HRG*s (blue) and *RPs* (green) is depicted for each histone dataset and condition. Analysis of the hypoxia responsive genes (*HRG*s) (n=49) and cytosolic ribosomal protein genes (n=223). Each row in the heat map represents an individual gene; gene order is the same in all plots. **b,c** Box plots of log_2_ fold change assayed for (**b**) short (2 h) and (**c**) prolonged (9 h) hypoxic stress compared to control (2HS/2NS, 9HS/9NS). Data for *ADH1* is indicated as a red dot and that of *RPL37B* is indicated as a blue dot. Range of dataset and upper and lower quantiles are depicted within each boxplot.

**Supplemental Figure 4.**
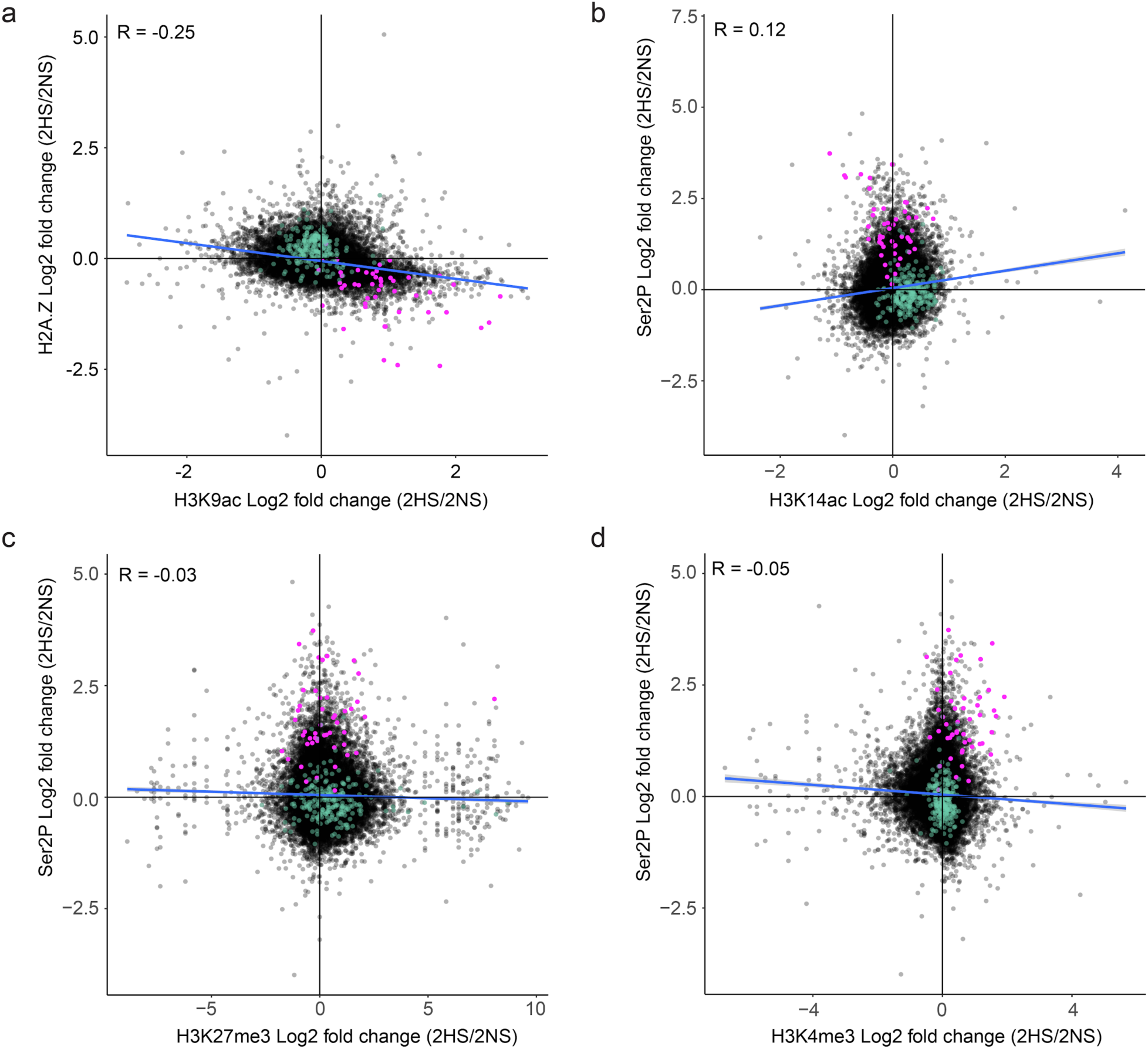
Evidence of cooperative association between H2A.Z eviction and H3K9ac and the relation between Histone H3 modification and RNAPII engagement. Plots comparing genome-wide changes in histone modifications and RNAP-Ser2P on individual gene bodies in response to hypoxia. **a** Comparison of log_2_ fold change (2HS/2NS) values of H3K9ac and H2A.Z association on gene bodies. **b-d** Comparison of log_2_ fold change (2HS/2NS) values of (**b**) H3K14ac, (**c**) H3K27me3, and (**d**) H3K4me3 and Ser2P on gene bodies. In all panels, *HRG*s are plotted in pink and *RP*s in green; the Pearson correlation coefficient for all genes is shown and the slope depicted with a blue line.

**Supplemental Figure 5.**
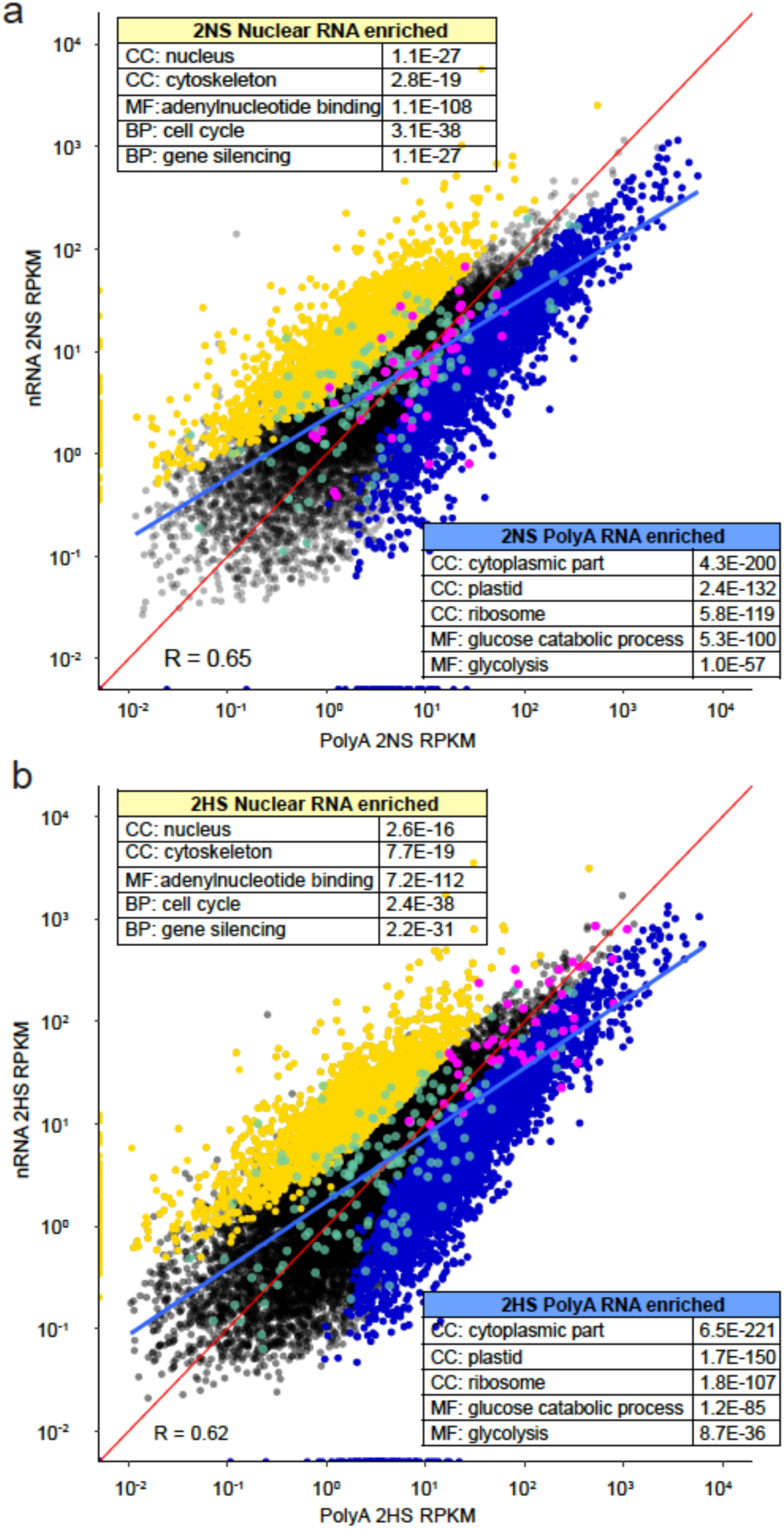
Genome-wide comparison of nRNA to polyA RNA demonstrates nuclear and cytoplasmic enrichment of transcripts under two conditions. **a, b** Scatterplot of normalized read values (per kilobase per million reads; RPKM) on genes in nRNA and polyA RNA populations at (**a**) 2NS and (**b**) 2HS. Transcripts enriched in nRNA (yellow; 2NS: 2,855, 2HS: 2,377) or polyA RNA (blue; 2NS: 3,139, 2HS: 4,029) are depicted (|log_2_ FC > 1|, p < 0.05). *HRG*s and *RP*s are also indicated in pink and green dots, respectively. The genome-wide Pearson correlation coefficient is shown and the slope depicted with a blue line. The red line is a slope of 1. Inserted tables describe representative Gene Ontology categories that are over-represented in nuclear RNA and polyA under the two conditions (data from Supplemental Table 2). Many of the same groups of cell component (CC) and molecular (MF) were over-represented in the nRNA and polyA RNA populations obtained in the same manner in rice (Reynoso et al., 2017).

**Supplemental Figure 6.**
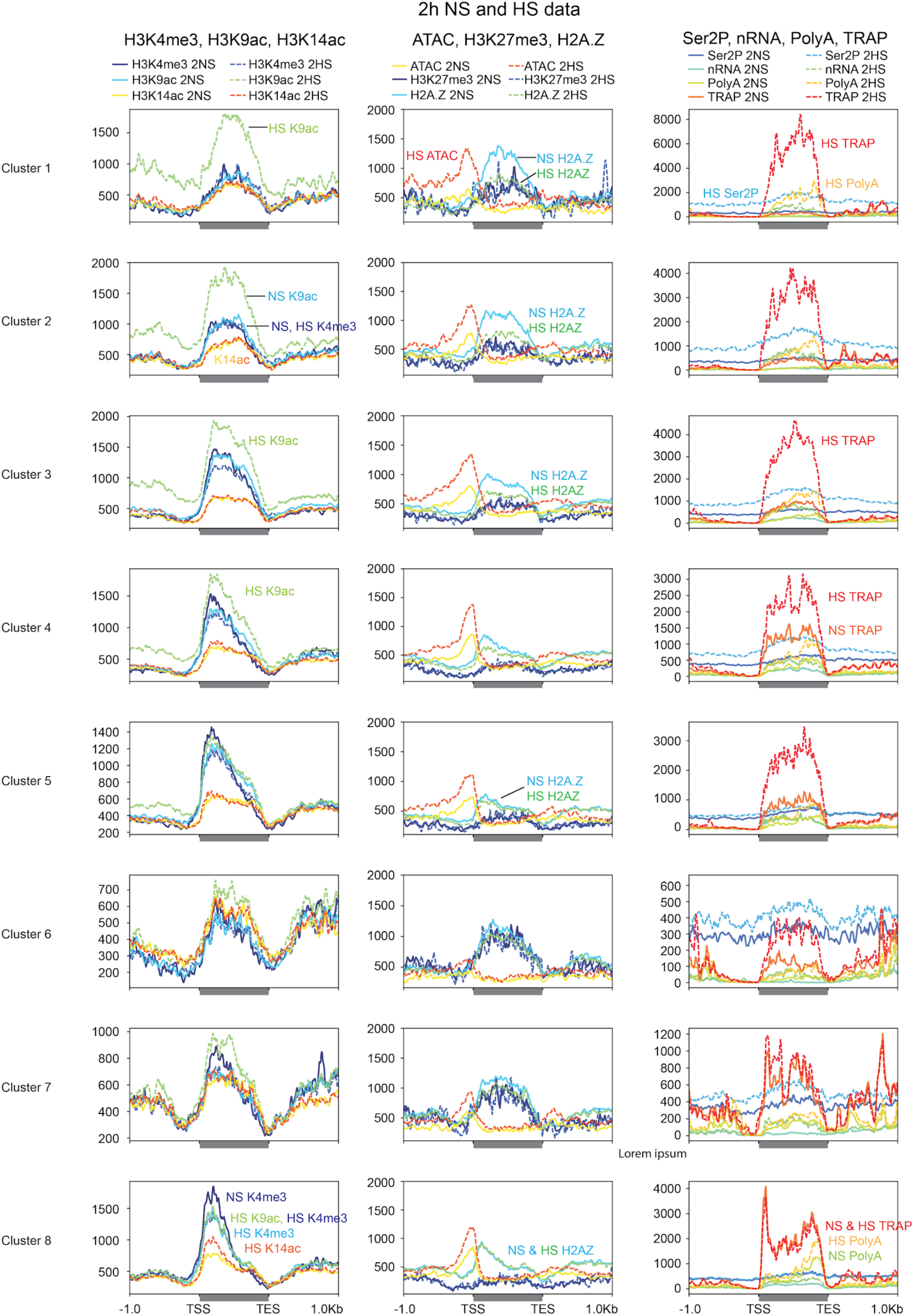
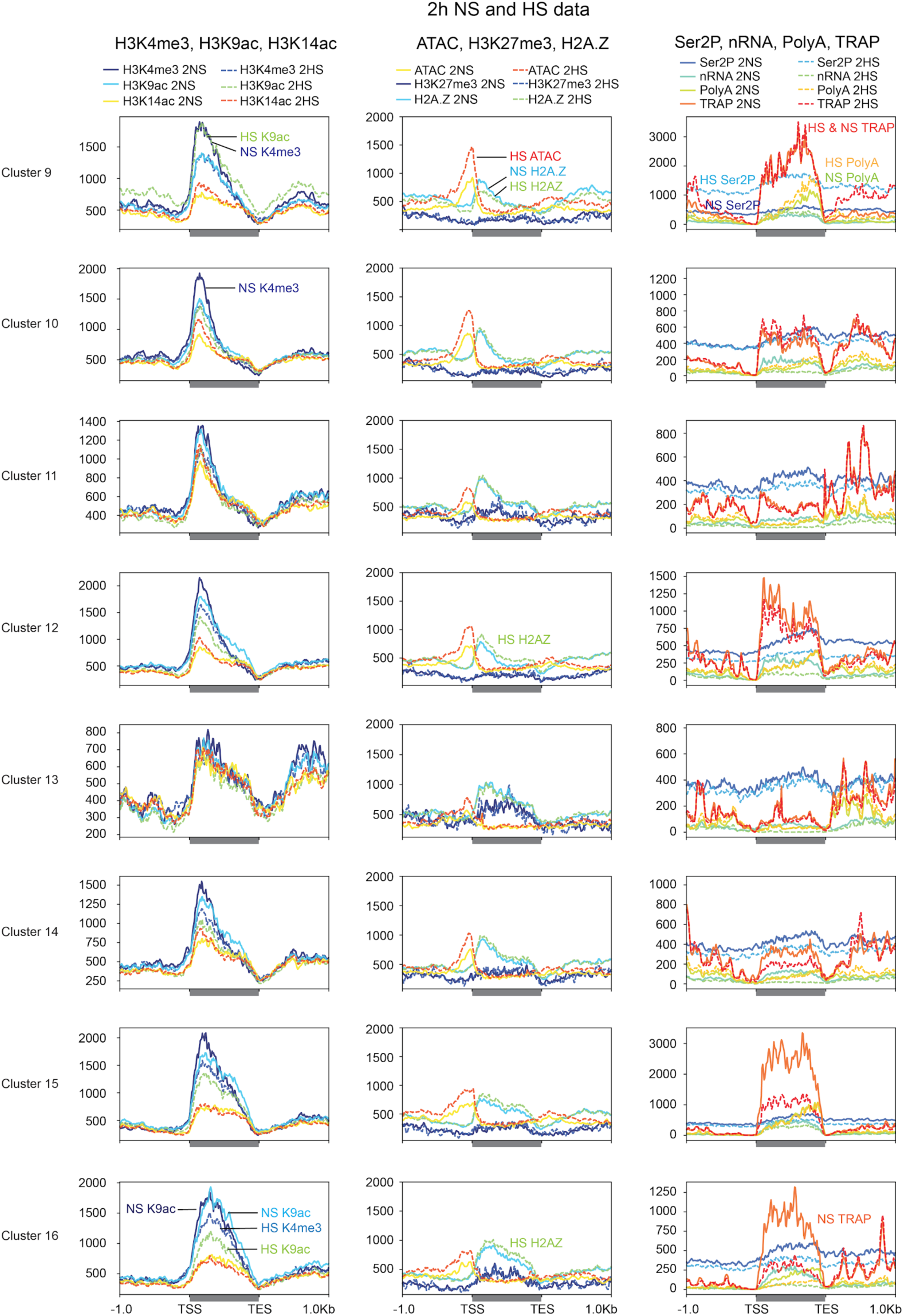
Distinctions in histones, RNAPII-Ser2P and RNA levels on genes that are co-regulated in response to hypoxic stress. Average signal of chromatin and RNA outputs for clustered genes identified from the 2HS comparison in Figure 4c. Hypoxic sample data are dashed lines. Signal values are plotted from 1 kb upstream of the TSS to 1 kb downstream of the TES. Signal scale of graphs may differ.

**Supplemental Figure 7.**
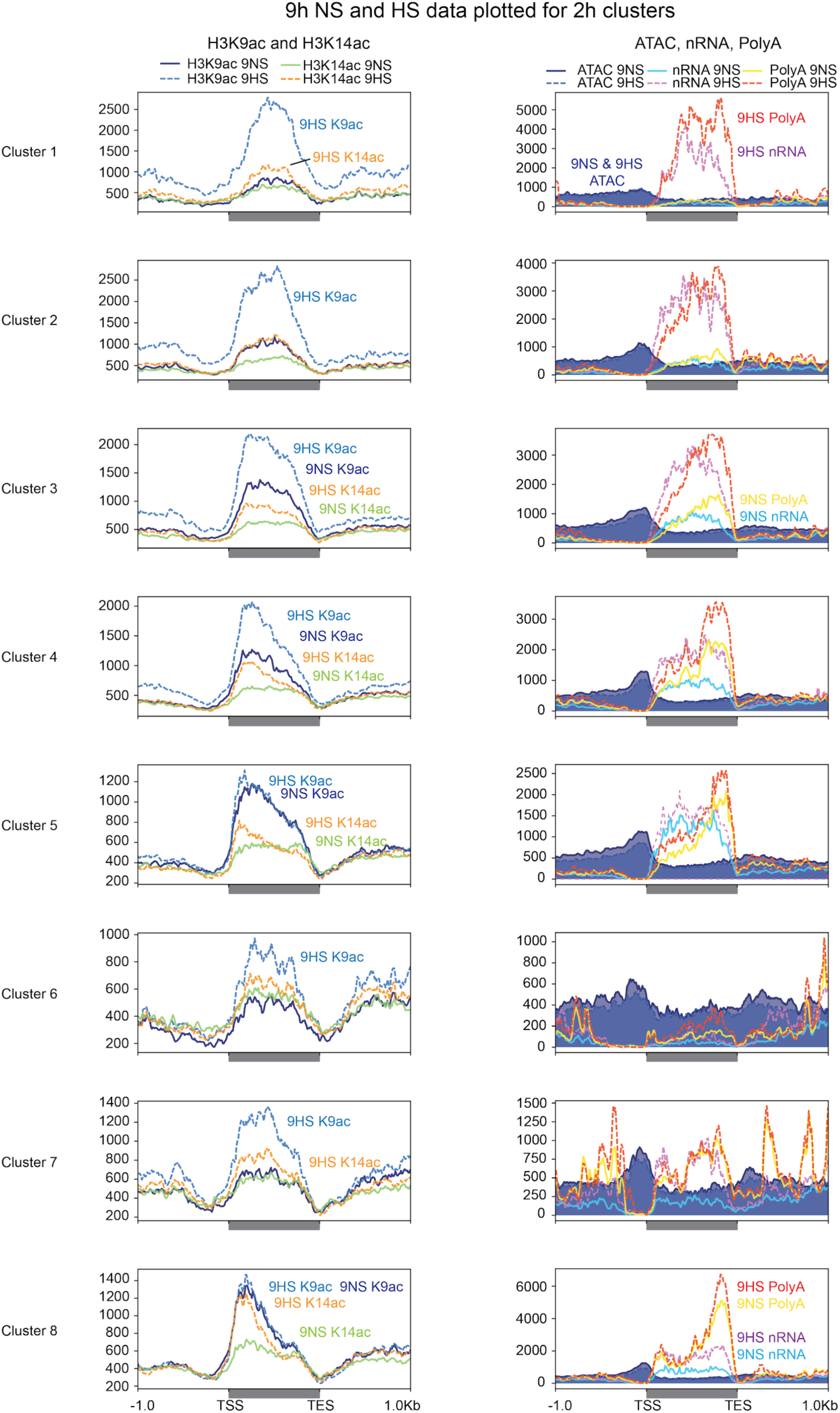
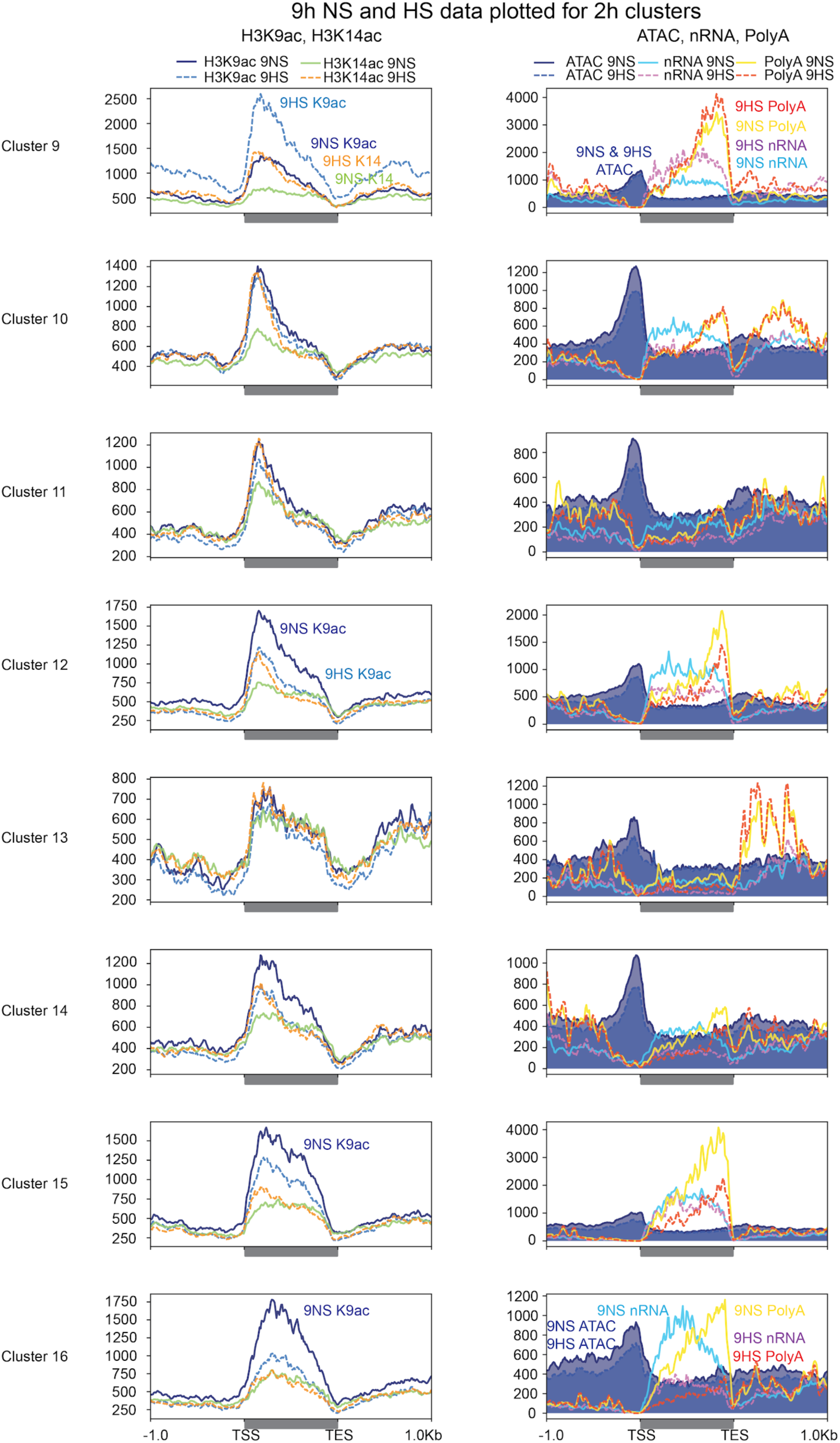
Progressive regulation of variations in gene activity in response to hypoxia. Average signal of chromatin and RNA outputs for clustered genes identified from the 9HS (9HS/9NS) comparison in Figure 4c. Plotting is as described for Supplemental Figure 6.

**Supplemental Figure 8.**
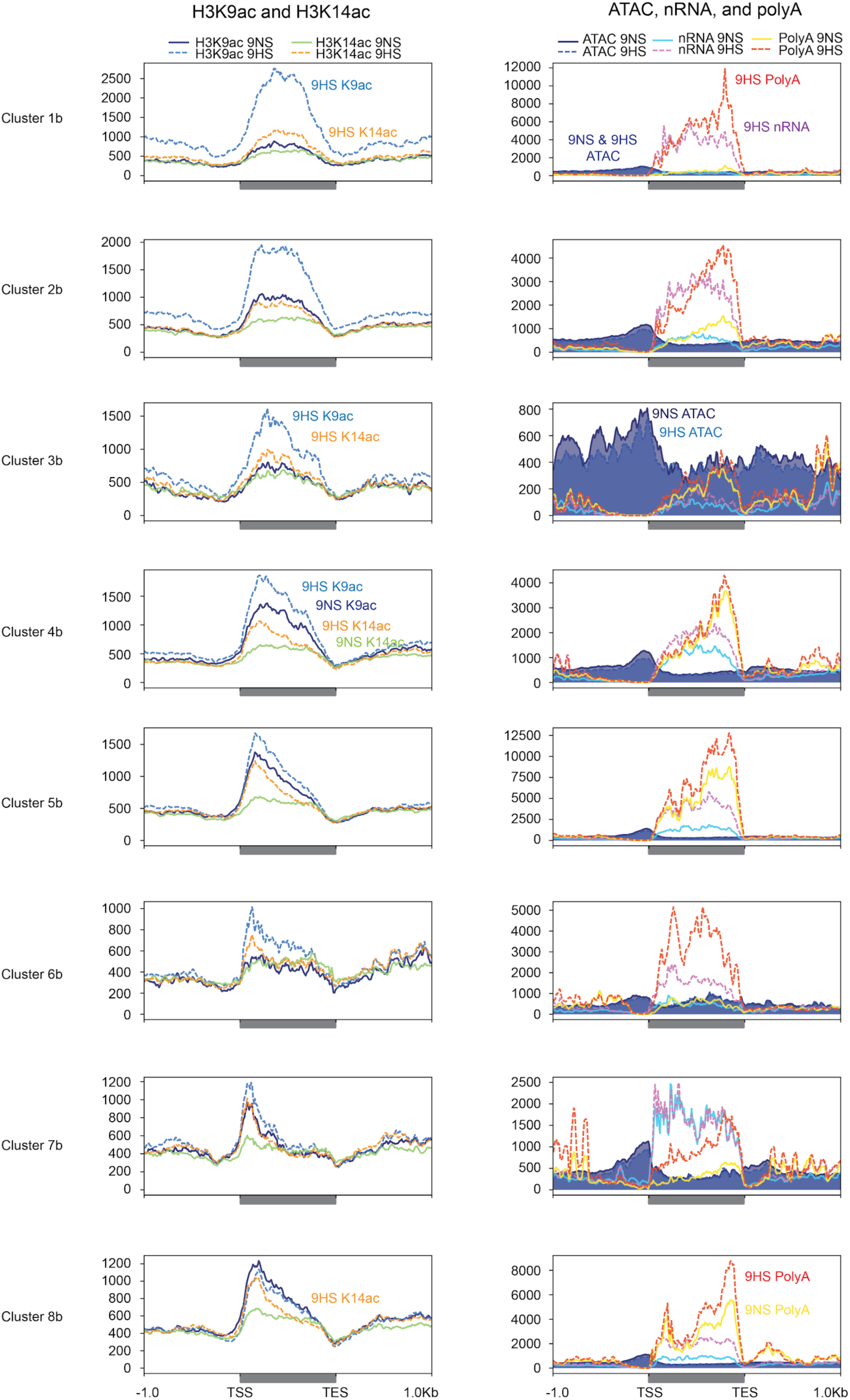
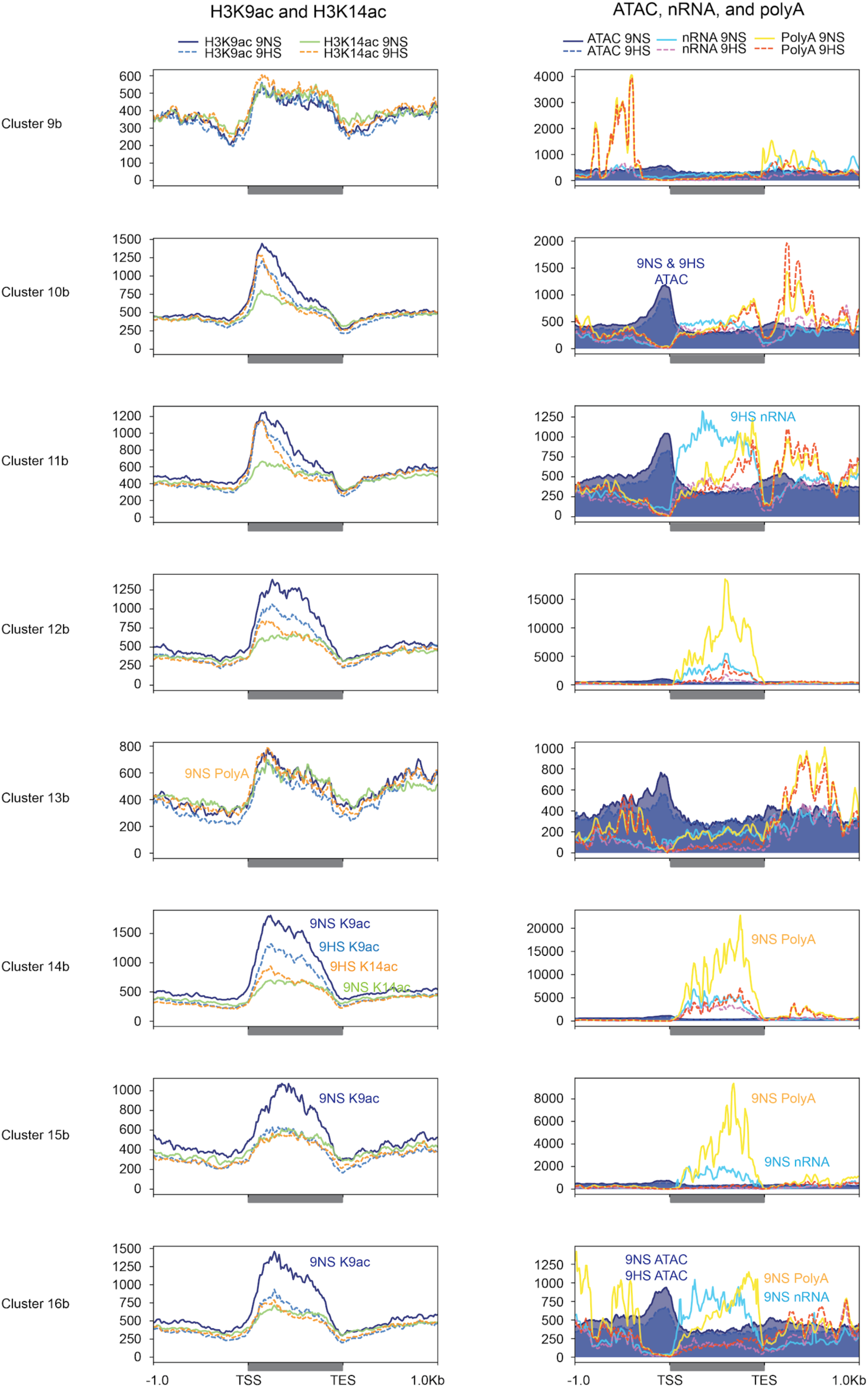
Histone and gene activity comparisons of genes co-regulated after 9 h of hypoxic stress. Average signal of chromatin and RNA data for co-regulated gene clusters identified from the 9 h hypoxic stress comparison (9HS/9NS) in Figure 6b. Plotting is as described for Supplemental Figure 6.

**Supplemental Figure 9.**
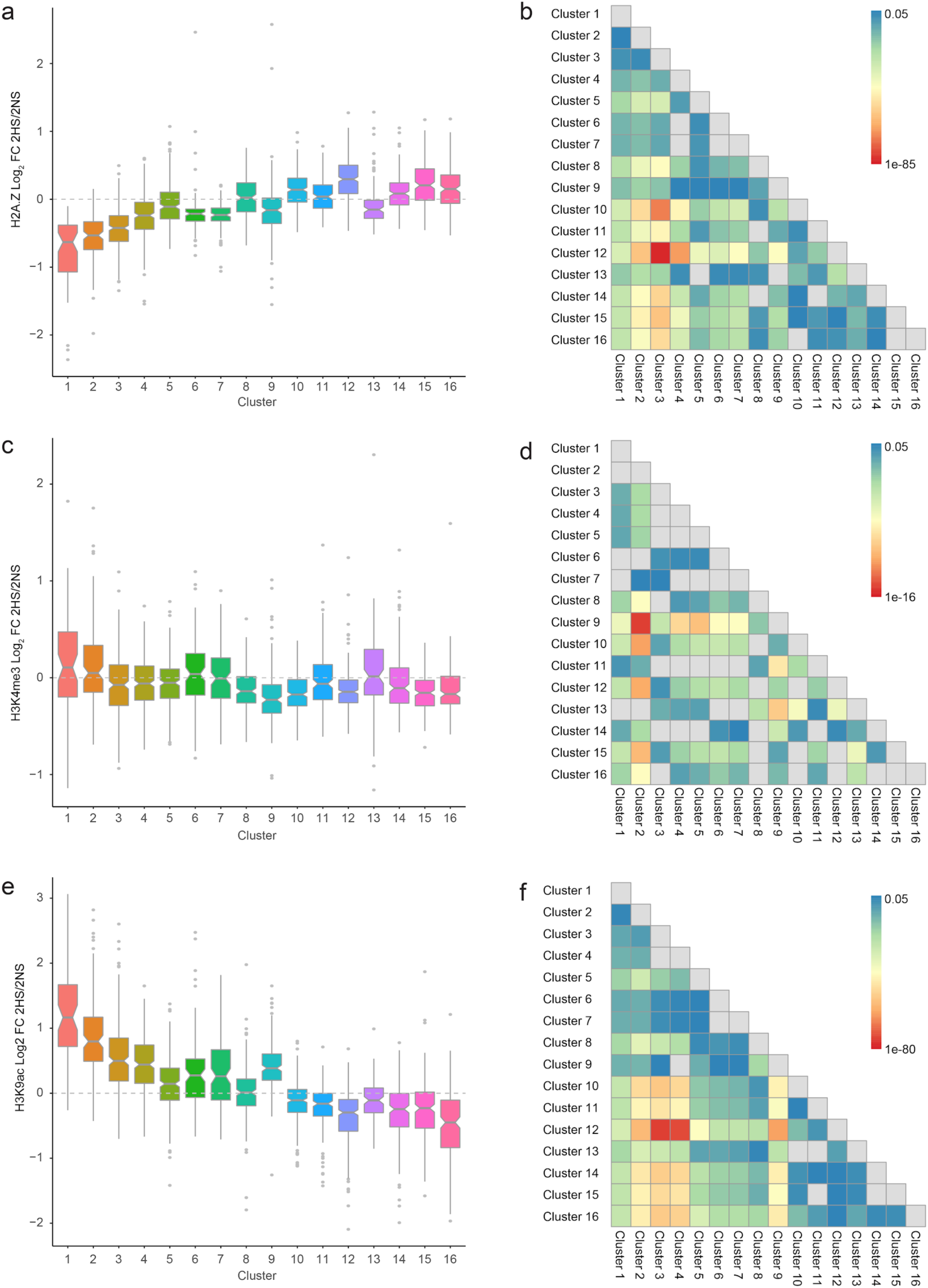
Quantitative analysis of dynamics in histone modifications of co-regulated gene clusters after 2 h of hypoxic stress. **a, c, e** Box plots of log_2_ fold change in response to 2 h hypoxic stress of H2A.Z, H3K4me3 and H3K9ac for the 16 clusters shown in Panels **b**, **d**, **f** provide Wilcoxon signed rank test values of gene clusters in an all by all matrix. Grey boxes denote no significant difference between clusters. Heatmap scale depicts significance values of p < 0.05.

**Supplemental Figure 10.**
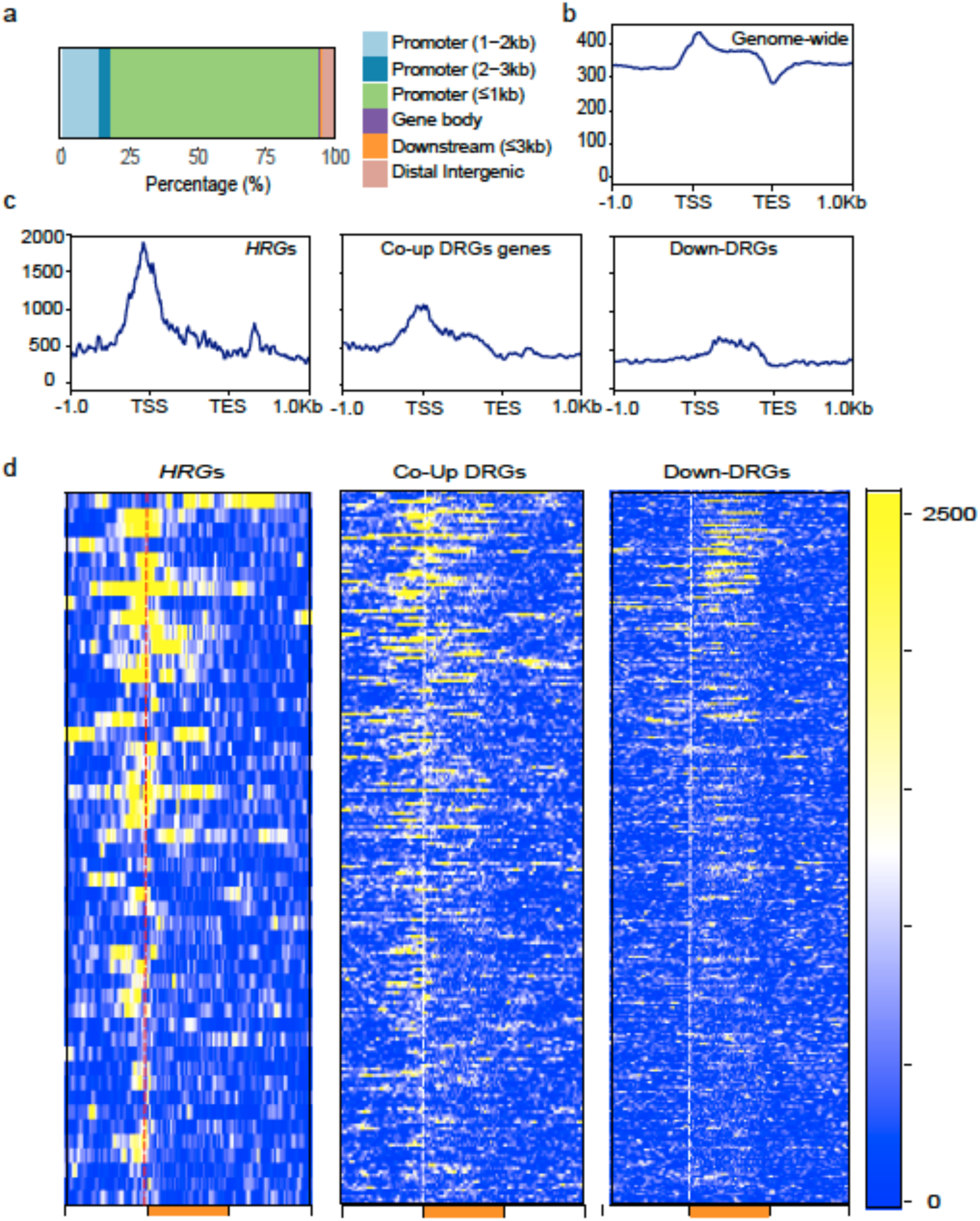
Genome-wide binding dynamics of HRE2 determined by ChIP-seq. **a** Distribution of 1,764 mapped HRE2 peaks on genomic features. **b, c** Average HRE2-ChIP read distribution in regions spanning 1 kb upstream to 1 kb downstream of gene features: **b** genome wide and **c** for the *HRG*s, Coordinately upregulated genes (Co-Up), and Down regulated genes (RGs). **d** Heat map of the HRE2-ChIP reads for genes in each group (*HRG*s) (n=49) and co-UP (n=213, and down-RGs (n=297). Each row in the heat map represents an individual gene; gene order is the same in all plots.

**Supplemental Figure 11.**
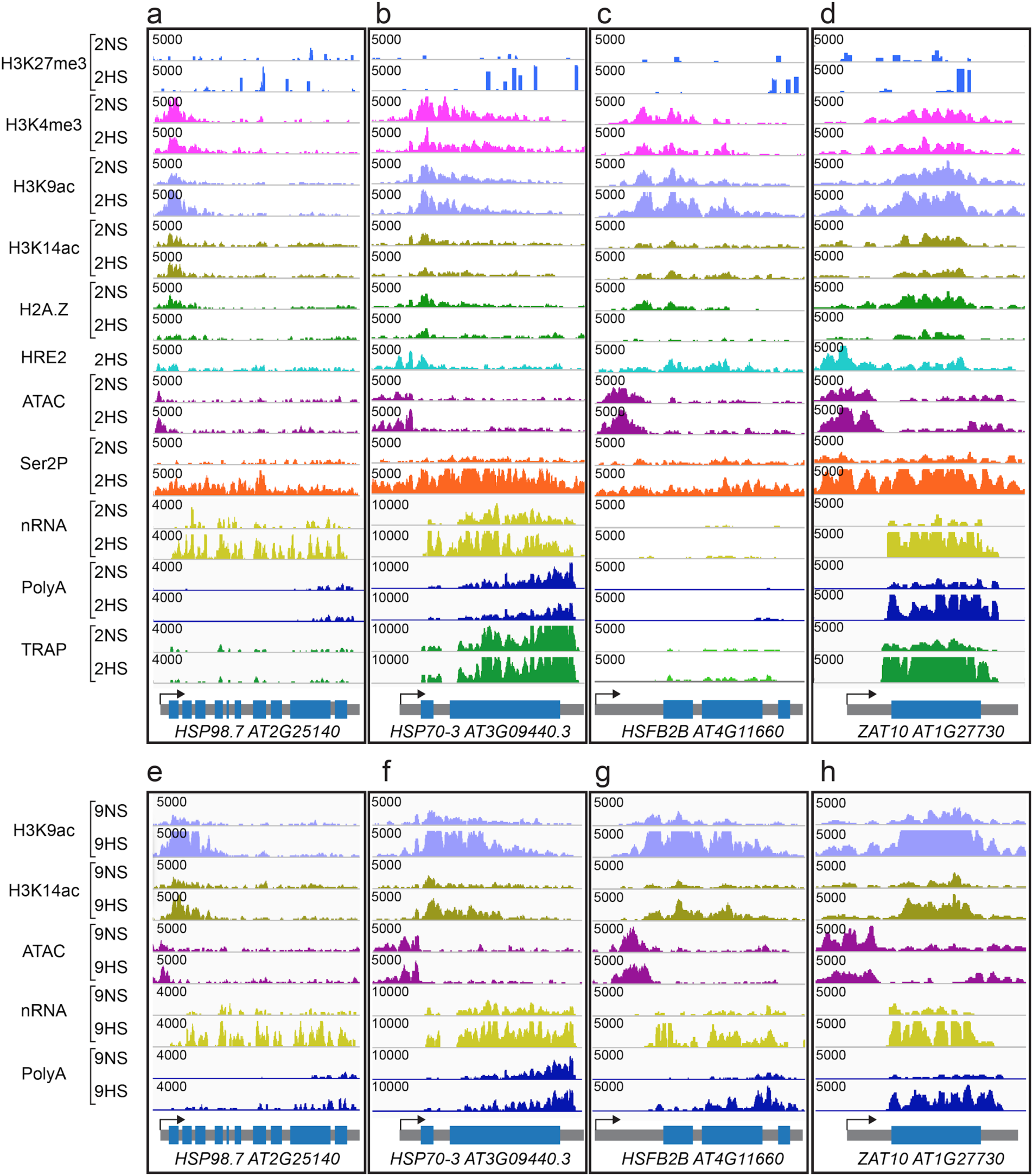
Genome browser view of normalized read coverage of histone, ATAC-seq, RNAPII-Ser2P, HRE2-chromatin immunopurification and RNA outputs for representative genes of cluster 9. **a-d** Normoxia (2NS) and 2 h hypoxic stress (2HS). **e-h** 9NS and 9HS samples. **a,e** *HEAT SHOCK PROTEIN (HSP) 98-7* **b,f** *HSP70-3,* **c,g** *HEAT SHOCK TRANSCRIPTION FACTOR B2B* and **d,h** *ZAT10.* The maximum read scale value used for chromatin based and RNA based outputs is provided. At bottom, the transcription unit is shown in grey with the transcription start site marked with an arrow.

**Supplemental Figure 12.**
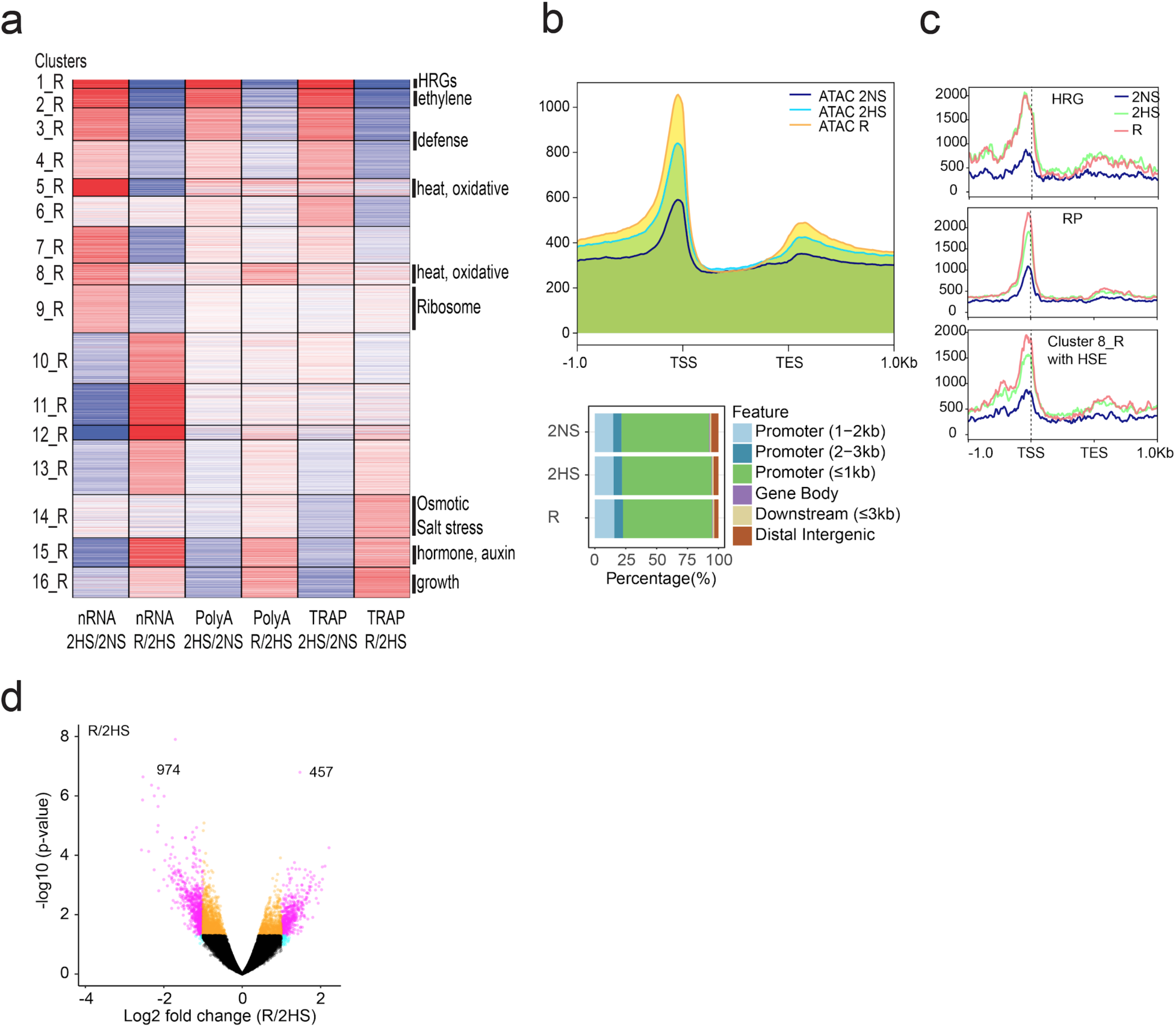
Reoxygenation following hypoxia promotes global dynamics in chromatin accessibility and RNA populations. **a** Heatmap showing similarly regulated genes based on dynamics in nRNA, polyA RNA, and TRAP in response to hypoxia (2HS/2NS) and reoxygenation (R/2HS). Partitioning around medoids (PAM) clustering of 4,784 differentially regulated genes; 16 clusters. Selected enriched Gene Ontology terms are shown at right (p adj. < 1.37E-06), |FC| > 1, FDR < 0.05. **a** number of clusters showed high nRNA at 2 HS that was not accompanied by high polyA or TRAP mRNA at 2HS clusters 5_R, 8_R, 10_R-13_R). Of these, cluster 8_R was the only with elevated polyA mRNA at R, suggesting these mRNAs accumulate to higher levels during reaeration. *HSP70-4* is a member of cluster 5_R. Both cluster 5_R and cluster 8_R genes are enriched for HSE elements within their promoters (cluster 5_R 85 of 123 genes have an HSE; cluster 8_R 81 of 153 genes have an HSE). Cluster 14_R is enriched for GOs associated with osmotic and salt stress. These stress-related mRNAs show preferential TRAP RNA accumulation at R, indicating a preferential translation at at this time point. **b** Average ATAC-seq read distribution of chromatin accessibility for all protein-coding genes at three timepoints and distribution of transposase hypersensitive sites on genomic features for each condition. **c** Dynamic chromatin accessibility in response to reaeration. 457 up- and 974 downregulated THSs. An increase in accessibility near the TSS was observed for the *RP*s but neither the *HRG*s nor the HSE-containing genes of cluster 8_R showed reduced accessibility within this time period. **d** Average signal of chromatin and RNA readouts for HRGs, RPs, and HSE containing cluster 8_R genes. Average signal is plotted from 1 kb upstream of the transcription start site (TSS) to 1 kb downstream of the transcript end site (TES). **d** Volcano plot of log_2_ fold change in THSs identifiable under 2HS and R conditions and the significance value of their difference. Genes indicated in pink meet the criteria of log_2_ fold change value > |1|, 0.05 < p. value.

**Supplemental Figure 13.**
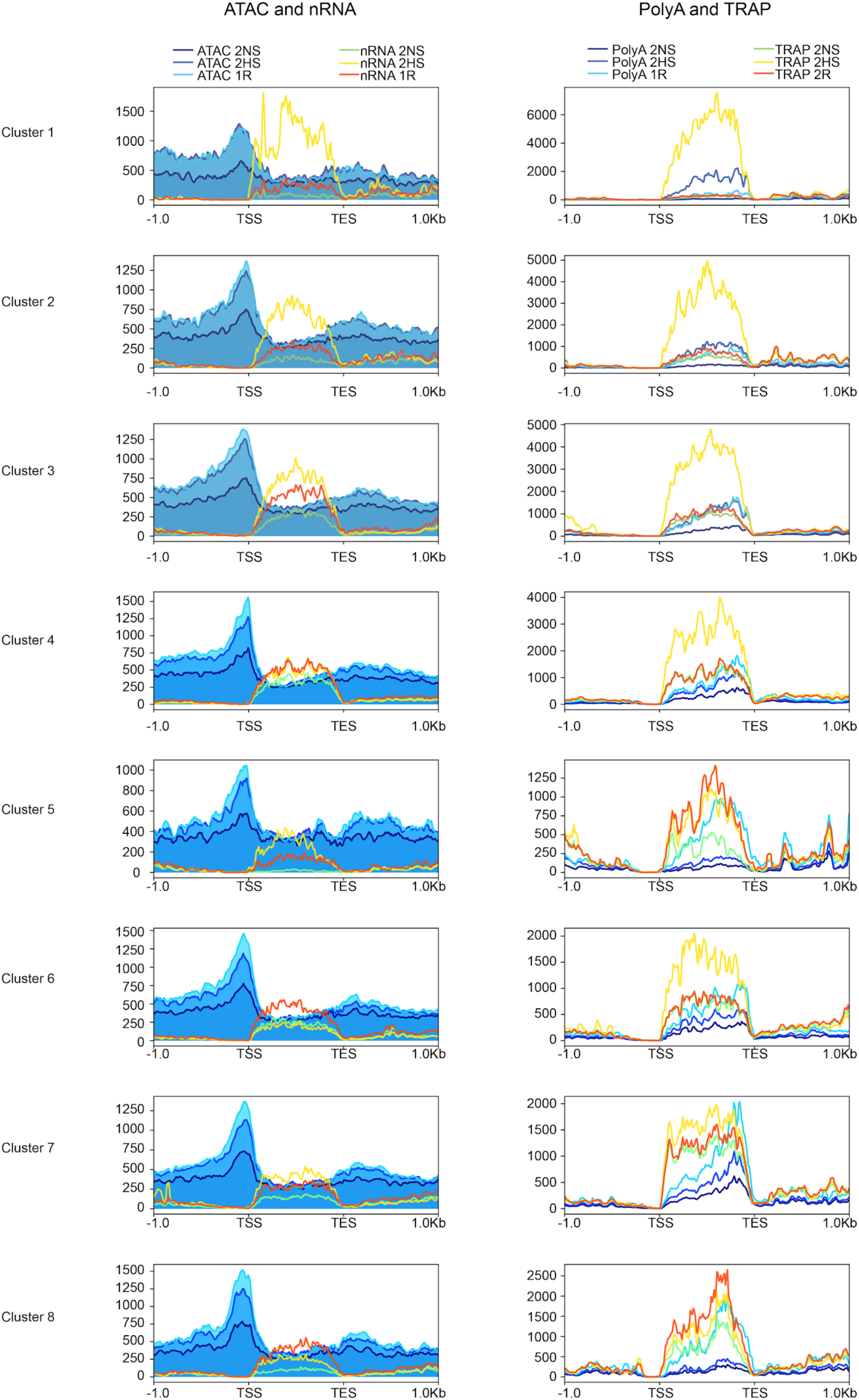
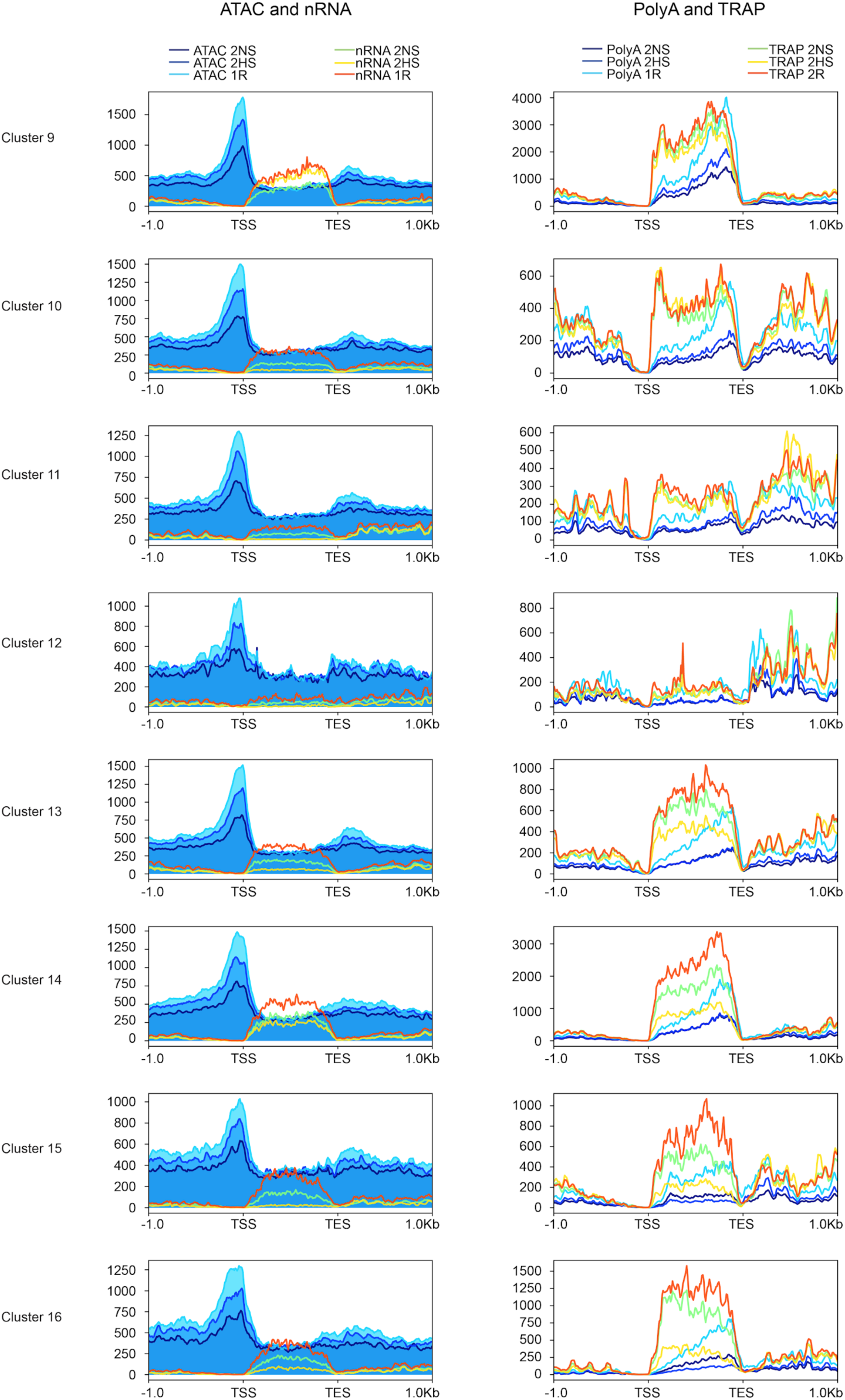
Regulation of variations in gene activity in response to hypoxia and reoxygenation. Average signal of chromatin and RNA outputs for clustered genes identified from the reoxygenation (2HS/2NS, R/2HS) comparison in Supplemental Figure 12. Plotting is as described for Supplemental Figure 6.

## References

Abbas, M., Berckhan, S., Rooney, D.J., Gibbs, D.J., Vicente Conde, J., Sousa Correia, C., Bassel, G.W., Marín-de la Rosa, N., León, J., Alabadí, D., Blázquez, M.A., and Holdsworth, M.J. (2015). Oxygen sensing coordinates photomorphogenesis to facilitate seedling survival. Curr. Biol. 25: 1483–1488.

Andrews, S. and Others (2010). FastQC: a quality control tool for high throughput sequence data.

Ansari, A. and Hampsey, M. (2005). A role for the CPF 3’-end processing machinery in RNAP II-dependent gene looping. Genes Dev. 19: 2969–2978.

Asensi-Fabado, M.-A., Amtmann, A., and Perrella, G. (2017). Plant responses to abiotic stress: The chromatin context of transcriptional regulation. Biochim. Biophys. Acta 1860: 106–122.

Bailey, T.L., Boden, M., Buske, F.A., Frith, M., Grant, C.E., Clementi, L., Ren, J., Li, W.W., and Noble, W.S. (2009). MEME SUITE: tools for motif discovery and searching. Nucleic Acids Res. 37: W202–8.

Banti, V., Loreti, E., Novi, G., Santaniello, A., Alpi, A., and Perata, P. (2008). Heat acclimation and cross-tolerance against anoxia in Arabidopsis. Plant Cell Environ. 31: 1029–1037.

Banti, V., Mafessoni, F., Loreti, E., Alpi, A., and Perata, P. (2010). The heat-inducible transcription factor HsfA2 enhances anoxia tolerance in Arabidopsis. Plant Physiol. 152: 1471–1483.

Batie, M., Frost, J., Frost, M., Wilson, J.W., Schofield, P., and Rocha, S. (2019). Hypoxia induces rapid changes to histone methylation and reprograms chromatin. Science 363: 1222–1226.

Baxter-Burrell, A., Yang, Z., Springer, P.S., and Bailey-Serres, J. (2002). RopGAP4-dependent Rop GTPase rheostat control of Arabidopsis oxygen deprivation tolerance. Science 296: 2026–2028.

Bernd Bischl, Michel Lang, Olaf Mersmann, Jörg Rahnenführer, and Claus Weihs (2015). BatchJobs and BatchExperiments&58; Abstraction Mechanisms for Using R in Batch Environments. J. Stat. Softw. 64: 1–25.

Blais, J.D., Filipenko, V., Bi, M., Harding, H.P., Ron, D., Koumenis, C., Wouters, B.G., and Bell, J.C. (2004). Activating transcription factor 4 is translationally regulated by hypoxic stress. Mol. Cell. Biol. 24: 7469–7482.

Branco-Price, C., Kaiser, K.A., Jang, C.J.H., Larive, C.K., and Bailey-Serres, J. (2008a). Selective mRNA translation coordinates energetic and metabolic adjustments to cellular oxygen deprivation and reoxygenation in Arabidopsis thaliana. Plant J. 56: 743–755.

Branco-Price, C., Kaiser, K.A., Jang, C.J.H., Larive, C.K., and Bailey-Serres, J. (2008b). Selective mRNA translation coordinates energetic and metabolic adjustments to cellular oxygen deprivation and reoxygenation in Arabidopsis thaliana. Plant J. 56: 743–755.

Branco-Price, C., Kawaguchi, R., Ferreira, R.B., and Bailey-Serres, J. (2005). Genome-wide analysis of transcript abundance and translation in Arabidopsis seedlings subjected to oxygen deprivation. Ann. Bot. 96: 647–660.

Buck, M.J., Raaijmakers, L.M., Ramakrishnan, S., Wang, D., Valiyaparambil, S., Liu, S., Nowak, N.J., and Pili, R. (2014). Alterations in chromatin accessibility and DNA methylation in clear cell renal cell carcinoma. Oncogene 33: 4961–4965.

Buenrostro, J.D., Giresi, P.G., Zaba, L.C., Chang, H.Y., and Greenleaf, W.J. (2013). Transposition of native chromatin for fast and sensitive epigenomic profiling of open chromatin, DNA-binding proteins and nucleosome position. Nat. Methods 10: 1213–1218.

Bui, L.T., Giuntoli, B., Kosmacz, M., Parlanti, S., and Licausi, F. (2015). Constitutively expressed ERF-VII transcription factors redundantly activate the core anaerobic response in Arabidopsis thaliana. Plant Sci. 236: 37–43.

Büttner, M. and Singh, K.B. (1997). Arabidopsis thaliana ethylene-responsive element binding protein (AtEBP), an ethylene-inducible, GCC box DNA-binding protein interacts with an ocs element binding protein. Proc. Natl. Acad. Sci. U. S. A. 94: 5961–5966.

Cao, J. et al. (2017). Comprehensive single-cell transcriptional profiling of a multicellular organism. Science 357: 661–667.

Chakraborty, A.A. et al. (2019). Histone demethylase KDM6A directly senses oxygen to control chromatin and cell fate. Science 363: 1217–1222.

Chang, R., Jang, C.J.H., Branco-Price, C., Nghiem, P., and Bailey-Serres, J. (2012). Transient MPK6 activation in response to oxygen deprivation and reoxygenation is mediated by mitochondria and aids seedling survival in Arabidopsis. Plant Mol. Biol. 78: 109–122.

Chervona, Y. and Costa, M. (2012). The control of histone methylation and gene expression by oxidative stress, hypoxia, and metals. Free Radic. Biol. Med. 53: 1041–1047.

Cortijo, S., Charoensawan, V., Brestovitsky, A., Buning, R., Ravarani, C., Rhodes, D., van Noort, J., Jaeger, K.E., and Wigge, P.A. (2017). Transcriptional Regulation of the Ambient Temperature Response by H2A.Z Nucleosomes and HSF1 Transcription Factors in Arabidopsis. Mol. Plant 10: 1258–1273.

Dai, X., Bai, Y., Zhao, L., Dou, X., Liu, Y., Wang, L., Li, Y., Li, W., Hui, Y., Huang, X., Wang, Z., and Qin, Y. (2017). H2A.Z Represses Gene Expression by Modulating Promoter Nucleosome Structure and Enhancer Histone Modifications in Arabidopsis. Mol. Plant 10: 1274–1292.

Deal, R.B. and Henikoff, S. (2010). A simple method for gene expression and chromatin profiling of individual cell types within a tissue. Dev. Cell 18: 1030–1040.

Ding, Y., Ndamukong, I., Xu, Z., Lapko, H., Fromm, M., and Avramova, Z. (2012). ATX1-generated H3K4me3 is required for efficient elongation of transcription, not initiation, at ATX1-regulated genes. PLoS Genet. 8: e1003111.

van Dongen, J.T. and Licausi, F. (2015). Oxygen sensing and signaling. Annu. Rev. Plant Biol. 66: 345–367.

Durinck, S. and Huber, W. biomaRt: Interface to BioMart databases (eg Ensembl, COSMIC, Wormbase and Gramene). R package version 2.

Eberharter, A. and Becker, P.B. (2002). Histone acetylation: a switch between repressive and permissive chromatin. Second in review series on chromatin dynamics. EMBO Rep. 3: 224–229.

Efroni, I., Han, S.-K., Kim, H.J., Wu, M.-F., Steiner, E., Birnbaum, K.D., Hong, J.C., Eshed, Y., and Wagner, D. (2013). Regulation of leaf maturation by chromatin-mediated modulation of cytokinin responses. Dev. Cell 24: 438–445.

ENCODE Project Consortium (2012). An integrated encyclopedia of DNA elements in the human genome. Nature 489: 57–74.

Eysholdt-Derzsó, E. and Sauter, M. (2017). Root Bending Is Antagonistically Affected by Hypoxia and ERF-Mediated Transcription via Auxin Signaling. Plant Physiol. 175: 412–423.

Fennoy, S.L., Nong, T., and Bailey-Serres, J. (1998). Transcriptional and post-transcriptional processes regulate gene expression in oxygen-deprived roots of maize. Plant J. 15: 727–735.

Gasch, P., Fundinger, M., Müller, J.T., Lee, T., Bailey-Serres, J., and Mustroph, A. (2015). Redundant ERF-VII transcription factors bind an evolutionarily-conserved cis-motif to regulate hypoxia-responsive gene expression in Arabidopsis. Plant Cell.

Gates, L.A., Foulds, C.E., and O’Malley, B.W. (2017). Histone Marks in the “Driver”s Seat’: Functional Roles in Steering the Transcription Cycle. Trends Biochem. Sci. 42: 977–989.

Gibbs, D.J. et al. (2014). Nitric oxide sensing in plants is mediated by proteolytic control of group VII ERF transcription factors. Mol. Cell 53: 369–379.

Gibbs, D.J., Lee, S.C., Isa, N.M., Gramuglia, S., Fukao, T., Bassel, G.W., Correia, C.S., Corbineau, F., Theodoulou, F.L., Bailey-Serres, J., and Holdsworth, M.J. (2011). Homeostatic response to hypoxia is regulated by the N-end rule pathway in plants. Nature 479: 415–418.

Girke, T. (2014). systemPipeR: NGS workflow and report generation environment. UC Riverside. https://github.com/tgirke/systemPipeR.

Giuntoli, B., Lee, S.C., Licausi, F., Kosmacz, M., Oosumi, T., van Dongen, J.T., Bailey-Serres, J., and Perata, P. (2014). A trihelix DNA binding protein counterbalances hypoxia-responsive transcriptional activation in Arabidopsis. PLoS Biol. 12: e1001950.

Giuntoli, B., Licausi, F., van Veen, H., and Perata, P. (2017). Functional Balancing of the Hypoxia Regulators RAP2.12 and HRA1 Takes Place in vivo in Arabidopsis thaliana Plants. Front. Plant Sci. 8: 591.

Giuntoli, B. and Perata, P. (2017). Group VII Ethylene Response Factors in Arabidopsis: regulation and physiological roles. Plant Physiol.

Gonzali, S., Loreti, E., Cardarelli, F., Novi, G., Parlanti, S., Pucciariello, C., Bassolino, L., Banti, V., Licausi, F., and Perata, P. (2015). Universal stress protein HRU1 mediates ROS homeostasis under anoxia. Nat Plants 1: 15151.

Gordon, A. and Hannon, G.J. (2010). Fastx-toolkit. FASTQ/A short-reads preprocessing tools (unpublished) http://hannonlab.cshl.edu/fastx_toolkit 5.

Hajheidari, M., Koncz, C., and Eick, D. (2013). Emerging roles for RNA polymerase II CTD in Arabidopsis. Trends Plant Sci. 18: 633–643.

Hattori, Y. et al. (2009). The ethylene response factors SNORKEL1 and SNORKEL2 allow rice to adapt to deep water. Nature 460: 1026–1030.

Heinz, S., Benner, C., Spann, N., Bertolino, E., Lin, Y.C., Laslo, P., Cheng, J.X., Murre, C., Singh, H., and Glass, C.K. (2010). Simple combinations of lineage-determining transcription factors prime cis-regulatory elements required for macrophage and B cell identities. Mol. Cell 38: 576–589.

Hetzel, J., Duttke, S.H., Benner, C., and Chory, J. (2016). Nascent RNA sequencing reveals distinct features in plant transcription. Proc. Natl. Acad. Sci. U. S. A. 113: 12316–12321.

Jonkers, I. and Lis, J.T. (2015). Getting up to speed with transcription elongation by RNA polymerase II. Nat. Rev. Mol. Cell Biol. 16: 167–177.

Juntawong, P., Girke, T., Bazin, J., and Bailey-Serres, J. (2014). Translational dynamics revealed by genome-wide profiling of ribosome footprints in Arabidopsis. Proc. Natl. Acad. Sci. U. S. A. 111: E203–12.

Karmodiya, K., Krebs, A.R., Oulad-Abdelghani, M., Kimura, H., and Tora, L. (2012). H3K9 and H3K14 acetylation co-occur at many gene regulatory elements, while H3K14ac marks a subset of inactive inducible promoters in mouse embryonic stem cells. BMC Genomics 13: 424.

Kim, D., Pertea, G., Trapnell, C., Pimentel, H., Kelley, R., and Salzberg, S.L. (2013). TopHat2: accurate alignment of transcriptomes in the presence of insertions, deletions and gene fusions. Genome Biol. 14: R36.

Kim, J.-M., To, T.K., Ishida, J., Matsui, A., Kimura, H., and Seki, M. (2012). Transition of chromatin status during the process of recovery from drought stress in Arabidopsis thaliana. Plant Cell Physiol. 53: 847–856.

Kim, J.-M., To, T.K., Ishida, J., Morosawa, T., Kawashima, M., Matsui, A., Toyoda, T., Kimura, H., Shinozaki, K., and Seki, M. (2008). Alterations of lysine modifications on the histone H3 N-tail under drought stress conditions in Arabidopsis thaliana. Plant Cell Physiol. 49: 1580–1588.

Kim, S.-H., Jeong, J.-W., Park, J.A., Lee, J.-W., Seo, J.H., Jung, B.-K., Bae, M.-K., and Kim, K.-W. (2007). Regulation of the HIF-1α stability by histone deacetylases. Oncol. Rep. 17: 647–651.

Kolde, R. (2012). Pheatmap: pretty heatmaps. R package version 61.

Koroleva, O.A., Calder, G., Pendle, A.F., Kim, S.H., Lewandowska, D., Simpson, C.G., Jones, I.M., Brown, J.W.S., and Shaw, P.J. (2009). Dynamic behavior of Arabidopsis eIF4A-III, putative core protein of exon junction complex: fast relocation to nucleolus and splicing speckles under hypoxia. Plant Cell 21: 1592–1606.

Kosmacz, M., Parlanti, S., Schwarzländer, M., Kragler, F., Licausi, F., and Van Dongen, J.T. (2015). The stability and nuclear localization of the transcription factor RAP2.12 are dynamically regulated by oxygen concentration. Plant Cell Environ. 38: 1094–1103.

Krijthe, J.H. (2015). Rtsne: T-distributed stochastic neighbor embedding using Barnes-Hut implementation. R package version 0. 13, URL https://github.com/jkrijthe/Rtsne.

Krzywinski, M., Schein, J., Birol, I., Connors, J., Gascoyne, R., Horsman, D., Jones, S.J., and Marra, M.A. (2009). Circos: an information aesthetic for comparative genomics. Genome Res. 19: 1639–1645.

Kumar, R., Ichihashi, Y., Kimura, S., Chitwood, D.H., Headland, L.R., Peng, J., Maloof, J.N., and Sinha, N.R. (2012). A High-Throughput Method for Illumina RNA-Seq Library Preparation. Front. Plant Sci. 3: 202.

Kumar, S.V. (2018). H2A.Z at the Core of Transcriptional Regulation in Plants. Mol. Plant 11: 1112–1114.

Kumar, S.V. and Wigge, P.A. (2010). H2A.Z-containing nucleosomes mediate the thermosensory response in Arabidopsis. Cell 140: 136–147.

Kwak, H. and Lis, J.T. (2013). Control of transcriptional elongation. Annu. Rev. Genet. 47: 483–508.

LaGory, E.L. and Giaccia, A.J. (2016). The ever-expanding role of HIF in tumour and stromal biology. Nat. Cell Biol. 18: 356–365.

Lawrence, M., Gentleman, R., and Carey, V. (2009). rtracklayer: an R package for interfacing with genome browsers. Bioinformatics 25: 1841–1842.

Lawrence, M., Huber, W., Pagès, H., Aboyoun, P., Carlson, M., Gentleman, R., Morgan, M.T., and Carey, V.J. (2013). Software for computing and annotating genomic ranges. PLoS Comput. Biol. 9: e1003118.

Licausi, F., Kosmacz, M., Weits, D.A., Giuntoli, B., Giorgi, F.M., Voesenek, L.A.C.J., Perata, P., and van Dongen, J.T. (2011). Oxygen sensing in plants is mediated by an N-end rule pathway for protein destabilization. Nature 479: 419–422.

Licausi, F., Van Dongen, J.T., Giuntoli, B., Novi, G., Santaniello, A., Geigenberger, P., and Perata, P. (2010). HRE1 and HRE2, two hypoxia-inducible ethylene response factors, affect anaerobic responses in Arabidopsis thaliana. Plant J. 62: 302–315.

Li, H. (2011). A statistical framework for SNP calling, mutation discovery, association mapping and population genetical parameter estimation from sequencing data. Bioinformatics 27: 2987–2993.

Lokdarshi, A., Conner, W.C., McClintock, C., Li, T., and Roberts, D.M. (2016). Arabidopsis CML38, a Calcium Sensor That Localizes to Ribonucleoprotein Complexes under Hypoxia Stress. Plant Physiol. 170: 1046–1059.

de Lorenzo, L., Sorenson, R., Bailey-Serres, J., and Hunt, A.G. (2017). Noncanonical Alternative Polyadenylation Contributes to Gene Regulation in Response to Hypoxia. Plant Cell 29: 1262–1277.

Loreti, E., Poggi, A., Novi, G., Alpi, A., and Perata, P. (2005). A genome-wide analysis of the effects of sucrose on gene expression in Arabidopsis seedlings under anoxia. Plant Physiol. 137: 1130–1138.

Lu, Z., Hofmeister, B.T., Vollmers, C., DuBois, R.M., and Schmitz, R.J. (2016). Combining ATAC-seq with nuclei sorting for discovery of cis-regulatory regions in plant genomes. Nucleic Acids Res.

Maechler, M. (2018). Finding Groups in Data’’: Cluster Analysis Extended Rousseeuw et. Documentation for software package. The Comprehensive R Archive Network (CRAN): Wien.

Maher, K.A. et al. (2018). Profiling of Accessible Chromatin Regions across Multiple Plant Species and Cell Types Reveals Common Gene Regulatory Principles and New Control Modules. Plant Cell 30: 15–36.

McCarthy, D.J., Chen, Y., and Smyth, G.K. (2012). Differential expression analysis of multifactor RNA-Seq experiments with respect to biological variation. Nucleic Acids Res. 40: 4288–4297.

Meyer, D., Dimitriadou, E., Hornik, K., Weingessel, A., and Leisch, F. (2015). e1071: Misc Functions of the Department of Statistics (e1071), TU Wien, 2014. R package version: 1–6.

Millar, A.H., Heazlewood, J.L., Giglione, C., Holdsworth, M.J., Bachmair, A., and Schulze, W.X. (2019). The Scope, Functions, and Dynamics of Posttranslational Protein Modifications. Annu. Rev. Plant Biol. 70: 119–151.

Milligan, L., Huynh-Thu, V.A., Delan-Forino, C., Tuck, A., Petfalski, E., Lombraña, R., Sanguinetti, G., Kudla, G., and Tollervey, D. (2016). Strand-specific, high-resolution mapping of modified RNA polymerase II. Mol. Syst. Biol. 12: 874.

Morgan, M., Anders, S., Lawrence, M., Aboyoun, P., Pagès, H., and Gentleman, R. (2009). ShortRead: a bioconductor package for input, quality assessment and exploration of high-throughput sequence data. Bioinformatics 25: 2607–2608.

Morgan, M., Carey, V., and Lawrence, M. (2014). BiocParallel: Bioconductor facilities for parallel evaluation. R package version 0. 4 1.

Morgan, M., Pages, H., Obenchain, V., and Hayden, N. (2016). Rsamtools: Binary alignment (BAM), FASTA, variant call (BCF), and tabix file import. R package version 1.

Mustroph, A., Lee, S.C., Oosumi, T., Zanetti, M.E., Yang, H., Ma, K., Yaghoubi-Masihi, A., Fukao, T., and Bailey-Serres, J. (2010). Cross-kingdom comparison of transcriptomic adjustments to low-oxygen stress highlights conserved and plant-specific responses. Plant Physiol. 152: 1484–1500.

Mustroph, A., Zanetti, M.E., Jang, C.J.H., Holtan, H.E., Repetti, P.P., Galbraith, D.W., Girke, T., and Bailey-Serres, J. (2009). Profiling translatomes of discrete cell populations resolves altered cellular priorities during hypoxia in Arabidopsis. Proc. Natl. Acad. Sci. U. S. A. 106: 18843–18848.

Neuwirth, E. (2011). RColorBrewer: colorbrewer palettes. R package version 1.

Niedojadło, J., Dełeńko, K., and Niedojadło, K. (2016). Regulation of poly(A) RNA retention in the nucleus as a survival strategy of plants during hypoxia. RNA Biol. 13: 531–543.

O’Malley, R.C., Huang, S.-S.C., Song, L., Lewsey, M.G., Bartlett, A., Nery, J.R., Galli, M., Gallavotti, A., and Ecker, J.R. (2016). Cistrome and Epicistrome Features Shape the Regulatory DNA Landscape. Cell 166: 1598.

Ozsolak, F., Poling, L.L., Wang, Z., Liu, H., Liu, X.S., Roeder, R.G., Zhang, X., Song, J.S., and Fisher, D.E. (2008). Chromatin structure analyses identify miRNA promoters. Genes Dev. 22: 3172–3183.

Paul, M.V., Iyer, S., Amerhauser, C., Lehmann, M., van Dongen, J.T., and Geigenberger, P. (2016). Oxygen Sensing via the Ethylene Response Transcription Factor RAP2.12 Affects Plant Metabolism and Performance under Both Normoxia and Hypoxia. Plant Physiol. 172: 141–153.

Phatnani, H.P. and Greenleaf, A.L. (2006). Phosphorylation and functions of the RNA polymerase II CTD. Genes Dev. 20: 2922–2936.

Pucciariello, C., Parlanti, S., Banti, V., Novi, G., and Perata, P. (2012). Reactive oxygen species-driven transcription in Arabidopsis under oxygen deprivation. Plant Physiol. 159: 184–196.

Quinlan, A.R. and Hall, I.M. (2010). BEDTools: a flexible suite of utilities for comparing genomic features. Bioinformatics 26: 841–842.

Ramírez, F., Ryan, D.P., Grüning, B., Bhardwaj, V., Kilpert, F., Richter, A.S., Heyne, S., Dündar, F., and Manke, T. (2016). deepTools2: a next generation web server for deep-sequencing data analysis. Nucleic Acids Res. 44: W160–5.

Reynoso, M., Pauluzzi, G., Kajala, K., Cabanlit, S., Velasco, J., Bazin, J., Deal, R., Sinha, N., Brady, S.M., and Bailey-Serres, J. (2017). Nuclear transcriptomes at high resolution using retooled INTACT. Plant Physiol.

Sato, F., Kitajima, S., and Koyama, T. (1996). Ethylene-induced gene expression of osmotin-like protein, a neutral isoform of tobacco PR-5, is mediated by the AGCCGCC eft-sequence. Plant Cell Physiol. 37: 249–255.

Sijacic, P., Bajic, M., McKinney, E.C., Meagher, R.B., and Deal, R.B. (2017). Chromatin accessibility changes between Arabidopsis stem cells and mesophyll cells illuminate cell type-specific transcription factor networks. bioRxiv: 213900.

Sorenson, R. and Bailey-Serres, J. (2014). Selective mRNA sequestration by OLIGOURIDYLATE-BINDING PROTEIN 1 contributes to translational control during hypoxia in Arabidopsis. Proc. Natl. Acad. Sci. U. S. A. 111: 2373–2378.

Sullivan, A.M. et al. (2014). Mapping and dynamics of regulatory DNA and transcription factor networks in A. thaliana. Cell Rep. 8: 2015–2030.

Sura, W., Kabza, M., Karlowski, W.M., Bieluszewski, T., Kus-Slowinska, M., Pawełoszek, Ł., Sadowski, J., and Ziolkowski, P.A. (2017). Dual Role of the Histone Variant H2A.Z in Transcriptional Regulation of Stress-Response Genes. Plant Cell 29: 791–807.

Talbert, P.B. and Henikoff, S. (2017). Histone variants on the move: substrates for chromatin dynamics. Nat. Rev. Mol. Cell Biol. 18: 115–126.

Torres, E.S. and Deal, R.B. (2018). The histone variant H2A. Z and chromatin remodeler BRAHMA act coordinately and antagonistically to regulate transcription and nucleosome dynamics in Arabidopsis. bioRxiv: 243790.

Townsley, B.T., Covington, M.F., Ichihashi, Y., Zumstein, K., and Sinha, N.R. (2015). BrAD-seq: Breath Adapter Directional sequencing: a streamlined, ultra-simple and fast library preparation protocol for strand specific mRNA library construction. Front. Plant Sci. 6: 366.

Van Veen, H., Vashisht, D., Akman, M., and Girke, T. (2016). Transcriptomes of eight Arabidopsis thaliana accessions reveal core conserved, genotype-and organ-specific responses to flooding stress. Plant.

Voesenek, L.A.C.J. and Bailey-Serres, J. (2013). Flooding tolerance: O2 sensing and survival strategies. Curr. Opin. Plant Biol. 16: 647–653.

Warnes, G.R., Bolker, B., Bonebakker, L., Gentleman, R., Huber, W., Liaw, A., Lumley, T., Maechler, M., Magnusson, A., Moeller, S., and Others (2009). gplots: Various R programming tools for plotting data. R package version 2: 1.

Watson, J.A., Watson, C.J., McCann, A., and Baugh, J. (2010). Epigenetics, the epicenter of the hypoxic response. Epigenetics 5: 293–296.

Weits, D.A., Giuntoli, B., Kosmacz, M., Parlanti, S., Hubberten, H.-M., Riegler, H., Hoefgen, R., Perata, P., van Dongen, J.T., and Licausi, F. (2014). Plant cysteine oxidases control the oxygen-dependent branch of the N-end-rule pathway. Nat. Commun. 5: 3425.

White, M.D. et al. (2017). Plant cysteine oxidases are dioxygenases that directly enable arginyl transferase-catalysed arginylation of N-end rule targets. Nat. Commun. 8: 14690.

Wickham, H. (2009). ggplot2: Elegant Graphics for Data Analysis (Springer Science & Business Media).

Wickham, H. and Francois, R. (2015). dplyr: A grammar of data manipulation. R package version 0. 4 3.

Xu, K., Xu, X., Fukao, T., Canlas, P., Maghirang-Rodriguez, R., Heuer, S., Ismail, A.M., Bailey-Serres, J., Ronald, P.C., and Mackill, D.J. (2006). Sub1A is an ethylene-response-factor-like gene that confers submergence tolerance to rice. Nature 442: 705–708.

Yang, C.-Y., Hsu, F.-C., Li, J.-P., Wang, N.-N., and Shih, M.-C. (2011). The AP2/ERF transcription factor AtERF73/HRE1 modulates ethylene responses during hypoxia in Arabidopsis. Plant Physiol. 156: 202–212.

Yang, W., Wightman, R., and Meyerowitz, E.M. (2017). Cell Cycle Control by Nuclear Sequestration of CDC20 and CDH1 mRNA in Plant Stem Cells. Mol. Cell 68: 1108–1119.e3.

Yeung, E. et al. (2018). A stress recovery signaling network for enhanced flooding tolerance in Arabidopsis thaliana. Proc. Natl. Acad. Sci. U. S. A. 115: E6085–E6094.

Young, R.M., Wang, S.-J., Gordan, J.D., Ji, X., Liebhaber, S.A., and Simon, M.C. (2008). Hypoxia-mediated selective mRNA translation by an internal ribosome entry site-independent mechanism. J. Biol. Chem. 283: 16309–16319.

Yu, G., Wang, L.-G., Han, Y., and He, Q.-Y. (2012). clusterProfiler: an R package for comparing biological themes among gene clusters. OMICS 16: 284–287.

Yu, G., Wang, L.-G., and He, Q.-Y. (2015). ChIPseeker: an R/Bioconductor package for ChIP peak annotation, comparison and visualization. Bioinformatics 31: 2382–2383.

Zanetti, M.E., Chang, I.-F., Gong, F., Galbraith, D.W., and Bailey-Serres, J. (2005). Immunopurification of polyribosomal complexes of Arabidopsis for global analysis of gene expression. Plant Physiol. 138: 624–635.

Zhou, J., Wang, X., He, K., Charron, J.-B.F., Elling, A.A., and Deng, X.W. (2010). Genome-wide profiling of histone H3 lysine 9 acetylation and dimethylation in Arabidopsis reveals correlation between multiple histone marks and gene expression. Plant Mol. Biol. 72: 585–595.

Zhu, J., Liu, M., Liu, X., and Dong, Z. (2018). RNA polymerase II activity revealed by GRO-seq and pNET-seq in Arabidopsis. Nat Plants.

Zid, B.M. and O’Shea, E.K. (2014). Promoter sequences direct cytoplasmic localization and translation of mRNAs during starvation in yeast. Nature 514: 117–121.

